# Discovery of frog virus 3 microRNAs and their roles in evasion of host antiviral responses

**DOI:** 10.1101/2021.09.17.460379

**Authors:** Lauren A. Todd, Barbara A. Katzenback

## Abstract

Frog virus 3 (FV3, genus *Ranavirus*) causes devastating disease in amphibian populations and is capable of subverting host immune responses. Evidence suggests that virus-encoded microRNAs (v-miRNAs) are implicated in host immunoevasion tactics. Thus, we sought to discover FV3-encoded v-miRNAs and to uncover their putative roles in immunoevasion. Small RNA libraries were generated from FV3-infected Xela DS2, a *Xenopus laevis* dorsal skin epithelial-like cell line, at 24- and 72-hours post-infection (hpi). We discovered 43 FV3 v-miRNAs and identified that 15 are upregulated at 24 hpi, while 18 are upregulated at 72 hpi. Target prediction analyses revealed that FV3 v-miRNAs target host genes involved in key antiviral signaling pathways, while gene ontology analyses suggest that FV3 v-miRNAs may broadly impact host cell function. This is the first study to experimentally detect mature v-miRNAs produced by FV3. Our findings highlight the possibility that ranaviral v-miRNAs facilitate immunoevasion of frog antiviral responses.

## Introduction

Ranaviruses are a group of large double-stranded DNA (dsDNA) viruses (family *Iridoviridae*) that cause devastating infections in amphibians and reptiles and represent a critical threat to amphibian biodiversity (Carey et al., 1999; Daszak et al., 1999; Peace et al., 2019; Price et al., 2014). Frog virus 3 (FV3) is the *Ranavirus* type species and is capable of infecting over 60 species of North American amphibians (Miller et al., 2011). Amphibian susceptibility to FV3 is influenced by various factors such as the environment, developmental stage, individual genotype, and host species (Brand et al., 2016; Gantress et al., 2003; Hoverman et al., 2010). In addition to infection of amphibians, FV3 can infect some reptile and fish species (Chinchar et al., 2009; McKenzie et al., 2019). The capacity for crossing species barriers suggests that FV3 is capable of evading host immune systems (Jacques et al., 2017). Indeed, initial studies have revealed that FV3 can shut down global host macromolecular synthesis, regardless of its ability to replicate (Goorha and Granoff, 1974; Maes and Granoff, 1967), and several viral genes encoded in the FV3 genome have been ascribed immunoevasive functions through the generation of FV3 gene knockout mutants (Chen et al., 2011; De Jesus Andino et al., 2015; Jacques et al., 2017). Additionally, we have previously observed that FV3 infection fails to elicit antiviral gene responses in susceptible frog skin epithelial-like cell lines, yet prior establishment of antiviral programs is protective against FV3 (Bui-Marinos et al., 2021), suggesting that FV3 is unable to break an established antiviral state but is able to prevent its establishment. Together, these observations suggest that FV3 is capable of immunoevasion and highlight the importance of broadening or understanding of how FV3 uses its genome to impact host immune responses.

The FV3 genome is 105.9 kilobases and encodes for at least 98 open reading frames (ORFs) which comprise 80% of the genome, while the remaining 20% of genomic space is occupied by intergenic regions harbouring regulatory elements such as untranslated regions (UTRs) (Tan et al., 2004). As with all viruses, FV3 interacts with host cellular machinery to drive viral replication. Studies suggest host transcription factors that are activated following FV3 infection [e.g. signal transducers and activators of transcription (STATs), interferon regulatory factors (IRFs), and nuclear factor kappa B (NF-κB)] can interact with putative promoter regions in the FV3 genome (Tian et al., 2021b). This suggests that host cells attempting to activate antiviral gene expression may inadvertently promote the transcription of viral genes, facilitating the production of viral molecules that may be involved in evading host immune responses and promoting viral replication.

While the functions of the majority of FV3 ORFs remain uncharacterized, a handful of studies have focused on understanding the role of FV3 ORFs in host-pathogen interactions and have implicated FV3 ORFs in immunoevasion. For example, FV3 encodes a caspase-like activation and recruitment domain (CARD) decoy-like molecule (vCARD) that interferes with interferon (IFN)-induced apoptosis in *Xenopus* hosts (De Jesus Andino et al., 2015; Jacques et al., 2017). It is believed that vCARD interacts with host immune signaling molecules that contain CARD domains such as RIG-I (retinoic acid-inducible gene 1), MDA5 (melanoma differentiation-associated protein 5), and MAVS (mitochondrial antiviral-signaling protein) to disrupt antiviral immunity (Besch et al., 2009; Jacques et al., 2017; Meylan et al., 2005). Additionally, FV3 encodes orthologs of several proteins known to play immunoevasion roles in other dsDNA viruses [reviewed in (Grayfer et al., 2015)] such as vIF-2α [viral homolog of eukaryotic initiation factor 2 alpha (eIF-2α); inhibits protein kinase R (PKR)-mediated eIF-2α phosphorylation], RNase III [blocks the interaction between viral double-stranded RNA (dsRNA) and PKR (Langland and Jacobs, 2002; Langland et al., 2006)], vβ-HSD [viral beta hydroxysteroid dehydrogenase; suppresses virus-induced cytopathic effects (Sun et al., 2006)], vTNFR [tumour necrosis factor (TNF) receptor; decoy molecule that blocks TNF], and DNA methyltransferase [blocks recognition by toll-like receptor 9 (TLR9) or cytoplasmic DNA sensors and prevents induction of IFN response (Krieg, 2002; Krug et al., 2004)]. Thus, it is apparent that FV3 ORFs play important roles in immunoevasion of host antiviral responses. Further characterization of FV3-encoded molecules will likely yield insight into additional mechanisms used by FV3 to evade host immune responses.

In the past two decades, microRNAs (miRNAs) have garnered significant attention in relation to host-pathogen interactions, and we previously discovered that frog miRNAs are implicated in host antiviral responses (Todd et al., accepted). miRNAs are small non-coding RNAs that function in post-transcriptional gene silencing through physical interactions with protein-coding gene transcripts. miRNAs can be produced by hosts and viruses, and viral miRNAs (v-miRNAs) have been detected in several dsDNA viruses afflicting humans including Epstein-Barr (EBV), herpes simplex virus type I (HSV-1), human cytomegalovirus (hCMV), and Kaposi sarcoma-associated herpesvirus (KSHV) (Kozomara et al., 2019). Due to the compact nature of viral genomes, v-miRNAs are not only encoded within intergenic regions but are often transcribed from genic regions (Cai et al., 2005; Cai et al., 2006; Pfeffer et al., 2005; Yuan et al., 2016). Functional interactions between v-miRNAs and their targets can occur in the 5’ UTR, coding sequence (cds), or 3’ UTR of the mRNA target (Grey et al., 2010; Lin and Ganem, 2011). v-miRNAs can target and regulate the expression of host proviral or antiviral genes or regulate endogenous viral gene expression (Duan et al., 2012; Enk et al., 2016; Grey et al., 2007; Lei et al., 2012; Lin et al., 2011; Lo et al., 2007; Lu et al., 2017; Scheel et al., 2017; Stern-Ginossar et al., 2007). By regulating host and viral gene expression, v-miRNAs have been shown to impact host responses to human viral pathogens including HSV-1, hCMV, EBV, and KSHV (Duan et al., 2012; Enk et al., 2016; Grey et al., 2007; Lei et al., 2012; Lin et al., 2011; Lo et al., 2007; Lu et al., 2017; Stern-Ginossar et al., 2007). Thus, v-miRNAs represent an attractive non-protein-coding evolutionary adaptation that can facilitate evasion of the host immune system.

Despite attracting less research attention than human viruses, a handful of iridovirids have also been found to encode v-miRNAs, including tiger frog virus (TFV), soft-shelled turtle iridovirus, and Singapore grouper iridovirus (SGIV) (Huang et al., 2009; Yan et al., 2011; Yuan et al., 2016). Interestingly, TFV-encoded v-miRNAs are thought to play a role in host-specificity (Yuan et al., 2016), and SGIV-encoded v-miRNAs have been shown to regulate viral gene expression by modulating major capsid protein expression which is thought to restrict early viral replication to prevent excessive cellular responses during infection (Yan et al., 2015). v-miRNA precursors have been computationally predicted from the FV3 genome (Li et al., 2008) and FV3 v-miRNAs have been predicted from transcriptionally active intergenic regions (Tian et al., 2021b), however mature FV3 v-miRNAs have yet to be detected experimentally. Since FV3 is known to use immunoevasion strategies to circumvent the host immune response, and other dsDNA viruses express v-miRNAs that target immune-related host genes to dampen host immune responses [reviewed in (Sorel and Dewals, 2016)], investigating the presence of FV3-encoded v-miRNAs may provide additional insight into how FV3 evades host immune responses throughout infection. Thus, the purpose of this study was to experimentally detect FV3-encoded v-miRNAs, and to assess their potential roles in regulating host antiviral gene expression and/or endogenous viral gene expression. As frog skin represents an important first line of defense against FV3 infection, we used a novel *Xenopus laevis* skin epithelial-like cell line, Xela DS2, that has been developed and characterized by our research group to address these questions.

## Materials and methods

### Cell culture medium

Xela complete medium was prepared as previously described (Bui-Marinos et al., 2020) and is composed of amphibian-adjusted Leibovitz’s L-15 (AL-15; Wisent, Mont-Saint-Hilaire, Canada) medium supplemented with 15% fetal bovine serum (FBS; lot # 234K18; VWR, Radnor, United States). Xela low serum medium is composed of AL-15 medium supplemented with 2% FBS. *Epithelioma Papulosum Cyprini* (EPC) complete medium is composed of Leibovitz’s L-15 medium supplemented with 10% FBS. EPC low serum medium is composed of L-15 medium supplemented with 2% FBS.

### Cell line maintenance

The Xela DS2 dorsal skin epithelial-like cell line was maintained in Xela complete medium as previously described (Bui-Marinos et al., 2020). Briefly, Xela DS2 cells were cultured at 26 °C in 75 cm^2^ plug-seal tissue-culture treated flasks (BioLite; Thermo Fisher Scientific, Waltham, United States) and confluent (~90%) cultures were split 1:4, every 3-4 days. EPC cells were maintained in EPC complete medium in 75 cm^2^ plug-seal tissue-culture treated flasks at 26 °C and were split 1:4 weekly. EPC cell lines are known to be contaminated with fathead minnow cells (Winton et al., 2010) but are still a valuable cell line for FV3 propagation.

### Viral titre determination for MOI calculations

FV3 (Granoff strain) was propagated in EPC cells as previously described (Bui-Marinos et al., 2021). A modified Kärber method (Kärber, 1931; Pham et al., 2011) was used to determine the tissue culture infectious dose in which 50% of cells are infected (TCID_50_/mL). Briefly, EPC cells were seeded in 96-well plates (BioBasic, Toronto, Canada) at a density of 100,000/well in EPC complete medium and adhered overnight at 26 °C. Cell culture medium was removed the next day and cells were treated with a ten-fold dilution series of FV3 stock diluted in EPC low serum medium. After incubation at 26 °C for seven days, EPC monolayers were visually examined for CPE to determine TCID_50_/mL values. Plaque forming unit (PFU)/mL values were determined by multiplying the calculated TCID_50_/mL values by 0.7 (Knudson and Tinsley, 1974). PFU/mL values associated with FV3 stocks were used to calculate multiplicity of infection (MOI) of 2 for FV3 infections.

### FV3 infection

Xela DS2 cells were seeded in 6-well plates at a density of 700,000 cells/well in 1 mL of Xela complete medium and incubated overnight at 26 °C. The next day, cell culture medium was removed and cell monolayers were treated with 2 mL of fresh Xela low serum medium (uninfected) or Xela low serum medium containing FV3 at an MOI of 2 (TCID_50_ of 2.0 × 10^6^ or 1.4 × 10^6^ PFUs; infected) for 2 h at 26 °C. Medium was removed and cells were washed three times with 1 mL of amphibian phosphate buffered saline (APBS; 8 g/L sodium chloride, 0.5 g/L potassium chloride, 2.68 g/L sodium phosphate dibasic heptahydrate, 0.24 g/L potassium phosphate monobasic) before the addition of 2 mL of fresh Xela low serum medium. Cells were incubated at 26 °C for the duration of the experiment.

### Sampling and total RNA isolation

The small RNA-seq data analyzed in this study originated from our previous paper examining changes in *X. laevis* miRNA expression in response to FV3 infection (Todd et al., accepted). Comprehensive details regarding sampling, RNA isolation, and small RNA library preparation and sequencing are described in the previous publication (Todd et al., accepted). In brief, total RNA was isolated from uninfected and FV3-infected Xela DS2 cells at 24 h post-infection (hpi) and 72 hpi using the Monarch Total RNA Minipreps kit (New England Biolabs, Ipswich, United States) and genomic DNA was removed using on-column DNase I digestion. Total RNA quantity and purity were assessed using a Cytation 5 Cell Imaging Multi-Mode Reader (Biotek, Winooski, United States) with the Take3 microvolume plate accessory (absorbance read at 230 nm, 260 nm, and 280 nm). RNA integrity was determined by bleach agarose gel electrophoresis (Aranda et al., 2012) and further analyzed on a Bioanalyzer 2100 using the RNA 6000 Nano LabChip Kit (Agilent Technologies Inc., Santa Clara, United States) by The Centre for Applied Genomics (TCAG) at The Hospital for Sick Children (Toronto, Canada) prior to small RNA sequencing.

### Reverse transcription PCR (RT-PCR)

To confirm infection of Xela DS2 cells, RT-PCR analysis was performed to detect viral gene expression. DNase I-treated total RNA (1 μg) was reverse-transcribed into cDNA using the BioBasic 5 × All-In-One RT PCR Master Mix. Resulting cDNA was diluted 1:8 and subjected to endpoint RT-PCR using Taq DNA Polymerase (GeneDirex Inc., Taoyuan, Taiwan) in a Mastercycler Nexus (Eppendorf) or Bio-Rad T100 thermocycler. PCR reaction conditions were as follows: 1 × GeneDirex buffer, 200 μM dNTPs, 0.2 μM forward and reverse primers, 0.625 U GeneDirex Taq, and 4 μL diluted cDNA. PCR cycling conditions for detection of FV3 *vCARD* were as follows: 95 °C for 5 min, 35 × (95 °C for 30 s; 58 °C for 30 s; 72 °C for 30 s), and 72 °C for 10 min. PCR cycling conditions for detection of FV3 *vPolIIα* were as follows: 95 °C for 5 min, 35 × (95 °C for 30 s; 58 °C for 30 s; 72 °C for 90 s), and 72 °C for 10 min. PCR cycling conditions for detection of FV3 *MCP* were as follows: 95 °C for 5 min, 35 × (95 °C for 45 s; 55 °C for 30 s; 72 °C for 60 s), and 72 °C for 10 min. *X. laevis actb* was detected as an internal control under the following cycling conditions: 95 °C for 5 min, 26 × (95 °C for 30 s, 55 °C for 30 s, 72 °C for 30 s), and 72 °C for 10 min. Primers used in this study are listed in **Table S1**.

### Small RNA sequencing (RNA-seq), read pre-processing, and read mapping

Small RNA libraries were generated from each RNA sample at TCAG (Toronto, Canada) using the NEBNext Multiplex Small RNA Library Prep Set for Illumina (New England Biolabs, Ipswich, United States) and sequenced on an Illumina HiSeq 2500 rapid-run flow cell to generate 50 nucleotide (nt) single-end reads by TCAG. As described in our previous paper (Todd et al., accepted), adaptor trimming and size filtering (17-30 nt) was performed on raw 50 nt single-end reads using Cutadapt (Martin, 2011) and quality control was performed using FastQC (Andrews, 2010). Processed reads that did not map to the *X. laevis* genome (Todd et al., accepted) were mapped to the FV3 genome (NC_005946.1) using Bowtie2 (Langmead and Salzberg, 2012) with the following parameters: single-end reads, write aligned and unaligned reads to separate files, allow one mismatch, output in SAM format. Cutadapt, FastQC, and Bowtie2 were executed using the Galaxy web platform (Afgan et al., 2016). FV3-aligned reads were retained for further analyses.

### Identification of FV3 v-miRNAs

As a positive control measure to validate our v-miRNA detection pipeline, we first used the pipeline to detect v-miRNAs from publicly available small RNA-seq datasets from EBV-positive human cell lines (C666-1, AKBM, and SNU-719) (Hooykaas et al., 2016). Small RNA reads were collapsed into unique reads (within each data set) with corresponding counts using miRDeep 2 software (v0.1.3; miRDeep2 mapper.pl script), which is the software that is most frequently used for *de novo* v-miRNA discovery from small RNA-seq data (Friedländer et al., 2012). Reads with a count < 10 were discarded for this purpose to improve the signal-to-noise ratio (Law et al., 2016). The collapsed reads were mapped to the human genome (hg19) using Bowtie2, and unaligned reads were subsequently mapped to the EBV genome (NC_009334.1) using Bowtie2. Files that were input into the miRDeep2.pl script included (1) collapsed EBV-mapped reads (at least 10 reads in a given library), (2) the EBV genome (NC_009334.1), (3) read alignments (converted from .sam to .arf format using the bwa_sam_converter.pl script), and (4) known v-miRNAs from miRBase (Kozomara et al., 2019). miRDeep2 analysis was executed using default parameters.

FV3 v-miRNAs were discovered from small RNA-seq libraries using miRDeep2 software as described above. Reads that mapped to the FV3 genome (and not the *X. laevis* genome) were collapsed into unique reads with associated counts according to treatment (e.g. 24 hpi replicates collapsed and 72 hpi replicates collapsed) using the miRDeep2 mapper.pl script, and reads with a count < 10 were discarded for this purpose. Files that were input into the miRDeep2.pl script included (1) collapsed FV3-mapped reads, (2) the FV3 genome (NC_005946.1), (3) read alignments (converted from .sam to .arf format using the bwa_sam_converter.pl script), and (4) a file containing known v-miRNAs from miRBase (Kozomara et al., 2019) and known *Iridovirid* v-miRNAs not in miRBase (Yan et al., 2016; Yan et al., 2011; Yuan et al., 2016; Zhang et al., 2014). Default mirDeep2.pl parameters were used. The resultant lists of FV3 v-miRNAs detected in the 24 hpi libraries and 72 hpi libraries were consolidated. Precursor secondary structures were generated using the RNAFold component of the ViennaRNA package v2.4.17 under default parameters (Lorenz et al., 2011).

### Quantification and differential expression analysis of novel FV3 v-miRNAs

FV3 v-miRNA read counts were generated using the miRDeep2 quantifier.pl script under default parameters. Collapsed FV3 reads, a fasta file containing FV3 v-miRNAs, and a fasta file containing FV3 pre-miRNAs served as input for the quantifier.pl script using default parameters. Resulting read count tables served as input for differential expression analyses (comparing FV3 v-miRNA levels at 24 hpi to their levels at 72 hpi) using DESeq2 (Love et al., 2014) and EdgeR (Robinson et al., 2010). DESeq2 parameters were as follows: select datasets per level, input data = count data, visualizing results = yes. EdgeR parameters were as follows: single count matrix, use factor information file = yes, use gene annotation = no, normalization method = TMM. FV3 v-miRNAs were considered differentially expressed if differential expression was identified by both programs (consensus) and FDR < 0.05. Heatmaps were generated using heatmap2 with the following parameters: enable data clustering = no, coloring groups = blue to white to red. DESeq2, EdgeR, and heatmap2 were run using the Galaxy web platform (Afgan et al., 2016).

### v-miRNA target prediction

v-miRNA target predictions were performed using RNAcalibrate and RNAhybrid v2.1.2 (Rehmsmeier et al., 2004). RNAcalibrate was used to estimate distribution parameters prior to running RNAhybrid (default parameters aside from strict seed pairing at bases 2-7 of the v-miRNA). FV3 v-miRNAs were examined for their likelihood of targeting *X. laevis* 3’ UTRs (retrieved from UCSC Table Browser using the Xenbase trackhub; genome.ucsc.edu; genome assembly v9.2) using RNAhybrid. RNAhybrid parameters were as follows: xi and theta values obtained from RNAcalibrate, maximum energy of −25 kcal/mol, maximum of one mismatch in internal loop, strict seed pairing at bases 2-7 of the v-miRNA, 0 mismatches in bulge loop, one hit per interaction, *p* < 0.01. Immune-related *X. laevis* targets of FV3 v-miRNAs were identified through a combination of manual search and cross-referencing The Gene Ontology Resource (The Gene Ontology Consortium, 2019) and InnateDB (Breuer et al., 2013). Interactions between FV3 v-miRNAs and endogenous FV3 transcripts were also predicted as described above using FV3 mRNA sequences retrieved from NCBI (Accession NC_005946.1).

### Gene ontology analysis

GO term enrichment analysis was performed using the PANTHER overrepresentation test to determine biological processes that are overrepresented in the list of *X. laevis* genes predicted to be targeted by FV3 v-miRNAs (Ashburner et al., 2000; Mi et al., 2019; The Gene Ontology Consortium, 2019). Test parameters were as follows: “GO biological process complete” annotation dataset, Fisher’s exact test, FDR < 0.05. GO analyses were performed at geneontology.org.

### Data availability

The small RNA-seq datasets generated by this study will be available in the NCBI Short Read Archive (SRA) repository (accession number pending manuscript acceptance).

## Results

### Validation of v-miRNA discovery pipeline

We sought to validate the ability of a bioinformatics analysis pipeline to accurately detect v-miRNAs and used EBV as an established system that is known to produce at least 44 v-miRNAs (Wang et al., 2018). Using publicly available small RNA-seq datasets from three EBV-positive human cell lines (Hooykaas et al., 2016), we were able to accurately detect 42 of the 44 known EBV v-miRNAs and discovered an additional 19 novel EBV v-miRNAs, some of which originate from the same precursor as a known v-miRNA and thus represent the other half of the miRNA:miRNA* duplex (**Table S2**). Thus, this pipeline appears to accurately detect v-miRNAs.

### FV3 encodes at least 43 v-miRNAs

In our previous study investigating the regulation of host miRNAs during FV3 infection (Todd et al., accepted), we noticed a substantial proportion of reads in the small RNA-seq libraries generated from FV3-infected samples (MOI of 2) that did not map to the *X. laevis* host genome. Since other iridovirids have been found to encode v-miRNAs, we mapped reads that did not align with the *X. laevis* genome to the FV3 genome using Bowtie2 (Langmead and Salzberg, 2012). The total number of FV3-mapped reads in each library ranged from 2.4–5.9 million (**Table 1**). The proportion of FV3-mapped reads in FV3-infected libraries increased over time from 9.3–14.5% (24 hpi) to 55.7–60.3% (72 hpi) (**Table 1**), which coincides with increases in viral transcript abundance (**Figure S1**) and viral titres (Todd et al., accepted). Conversely, a very small number of reads (0.004% or less) in libraries generated from uninfected cells mapped to the FV3 genome, which likely represent sequencing artifacts. These reads were discarded prior to analysis, as their source cannot be determined. Forty-three v-miRNAs, between 18 – 25 nt, which are produced from 33 pre-miRNAs (**Figure 1** and **Table S3**) were discovered using miRDeep2 software (Friedländer et al., 2012) (**Table 2**). FV3 v-miRNAs were mapped to the FV3 genome (**Figure 2**), revealing that they are encoded in various orientations relative to viral protein-coding genes. Nineteen FV3 v-miRNAs are produced from genic regions in the sense direction (i.e. same strand as protein-coding gene), 21 v-miRNAs are produced from genic regions in the antisense direction (i.e. strand opposite protein-coding gene), and three FV3 v-miRNAs are produced from intergenic regions (**Figure 2** and **Table 2**). Alignment of these v-miRNAs to *X. laevis* miRNAs in miRBase and those previously identified by our group (Todd et al., accepted) revealed no seed sequence conservation with *X. laevis* miRNAs (**Table 2**). Alignment of FV3 v-miRNAs to v-miRNAs encoded by other iridovirids (SGIV, TFV, megalocytivirus, and infectious kidney necrosis virus) and non-iridovirid v-miRNAs in miRBase (Kozomara et al., 2019) revealed seed sequence conservation with v-miRNAs from one iridovirid (TFV) as well as non-iridovirid dsDNA viruses including Marek’s disease virus (MDV), Rhesus lymphocryptovirus (rLCV), Gorilla gorilla polyomavirus 1 (gggpv1), and EBV (**Table 2**).

**Figure 1.**
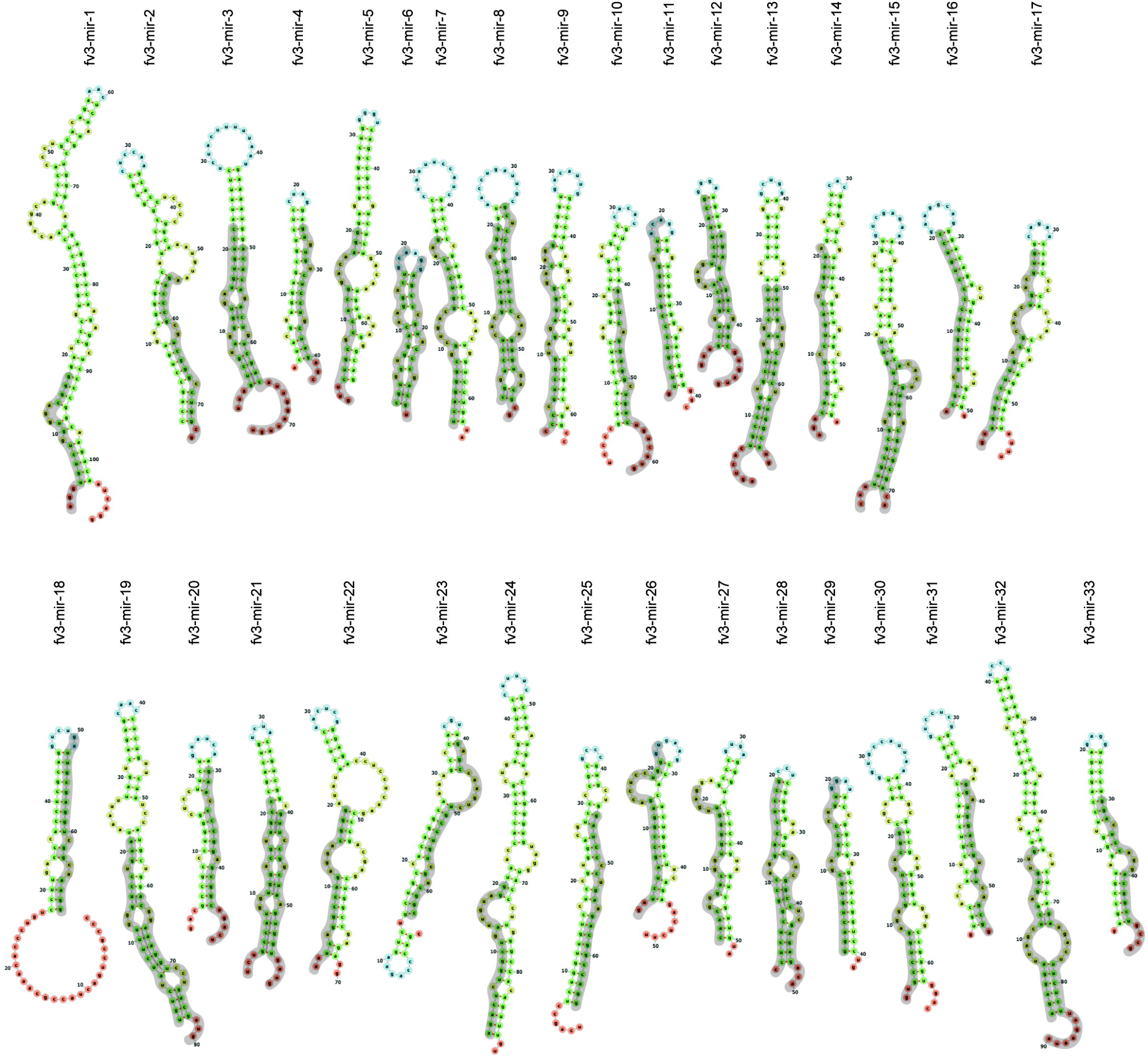
Structure of FV3 pre-miRNAs and mature v-miRNAs. Pre-miRNAs identified by miRDeep2 were input into the *forna* web server (rna.tbi.univie.ac.at/forna) to determine hairpin structures. Mature v-miRNA locations in pre-miRNA hairpin structures are highlighted in grey. Terminal loops are represented in blue, internal loops are represented in yellow, helices are represented in green, and dangling ends/terminal mismatches are represented in orange.

**Figure 2.**
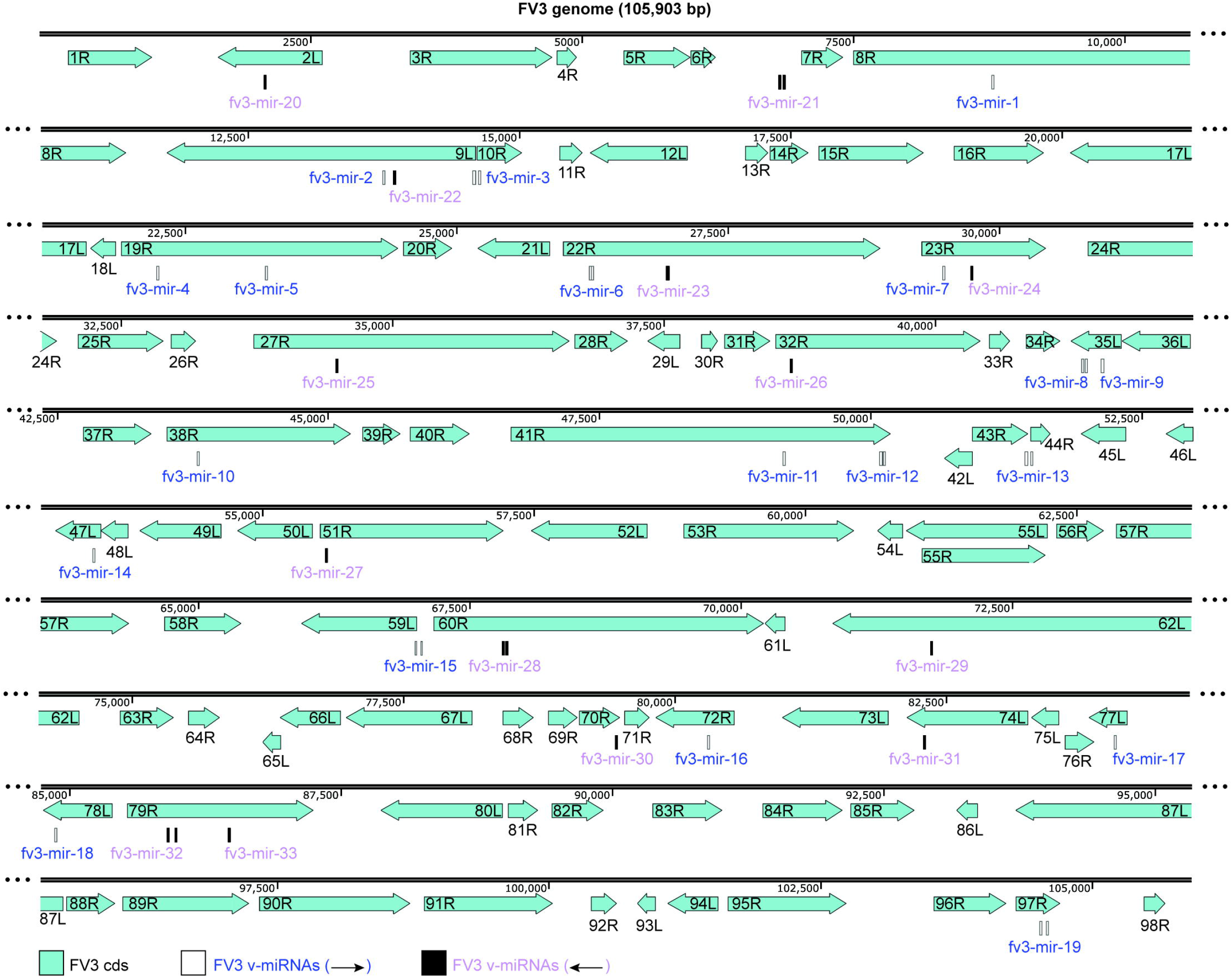
Schematic of FV3 v-miRNAs aligned to the FV3 genome. FV3 coding sequences (cds) are represented by teal arrows, and corresponding open reading frame numbers are indicated. Mature FV3 v-miRNAs encoded on the plus strand are depicted by white rectangles (labeled with blue text), while mature FV3 v-miRNAs encoded on the minus strand are depicted by black rectangles (labeled with purple text). The 5p and 3p mature v-miRNAs are labeled with the name of their respective pre-miRNA due to space constraints. bp, base-pair.

**Table 1.** Number of FV3-mapped reads in small RNA-seq libraries.

| Small RNA-seq library | Total reads (17-30 nt) | FV3-mapped reads (17-30 nt) |
| --- | --- | --- |
| Uninfected 24 hpi #1 | 3,902,750 | 156 (0.004%)* |
| Uninfected 24 hpi #2 | 3,704,214 | 70 (0.002%)* |
| Uninfected 24 hpi #3 | 3,831,323 | 61 (0.002%)* |
| Uninfected 72 hpi #1 | 3,134,642 | 116 (0.004%)* |
| Uninfected 72 hpi #2 | 2,415,696 | 95 (0.004%)* |
| Uninfected 72 hpi #3 | 3,654,713 | 67 (0.002%)* |
| FV3-infected 24 hpi #1 | 5,329,573 | 495,978 (9.306%) |
| FV3-infected 24 hpi #2 | 2,681,465 | 391,085 (14.585%) |
| FV3-infected 24 hpi #3 | 3,162,325 | 296,877 (9.388%) |
| FV3-infected 72 hpi #1 | 5,906,452 | 3,288,433 (55.675%) |
| FV3-infected 72 hpi #2 | 5,440,609 | 3,284,168 (60.364%) |
| FV3-infected 72 hpi #3 | 5,213,772 | 3,035,202 (58.215%) |

\*A very small number of reads in uninfected samples mapped to the FV3 genome. These reads were excluded from all libraries prior to analysis as their source is unable to be accurately determined.

**Table 2.**
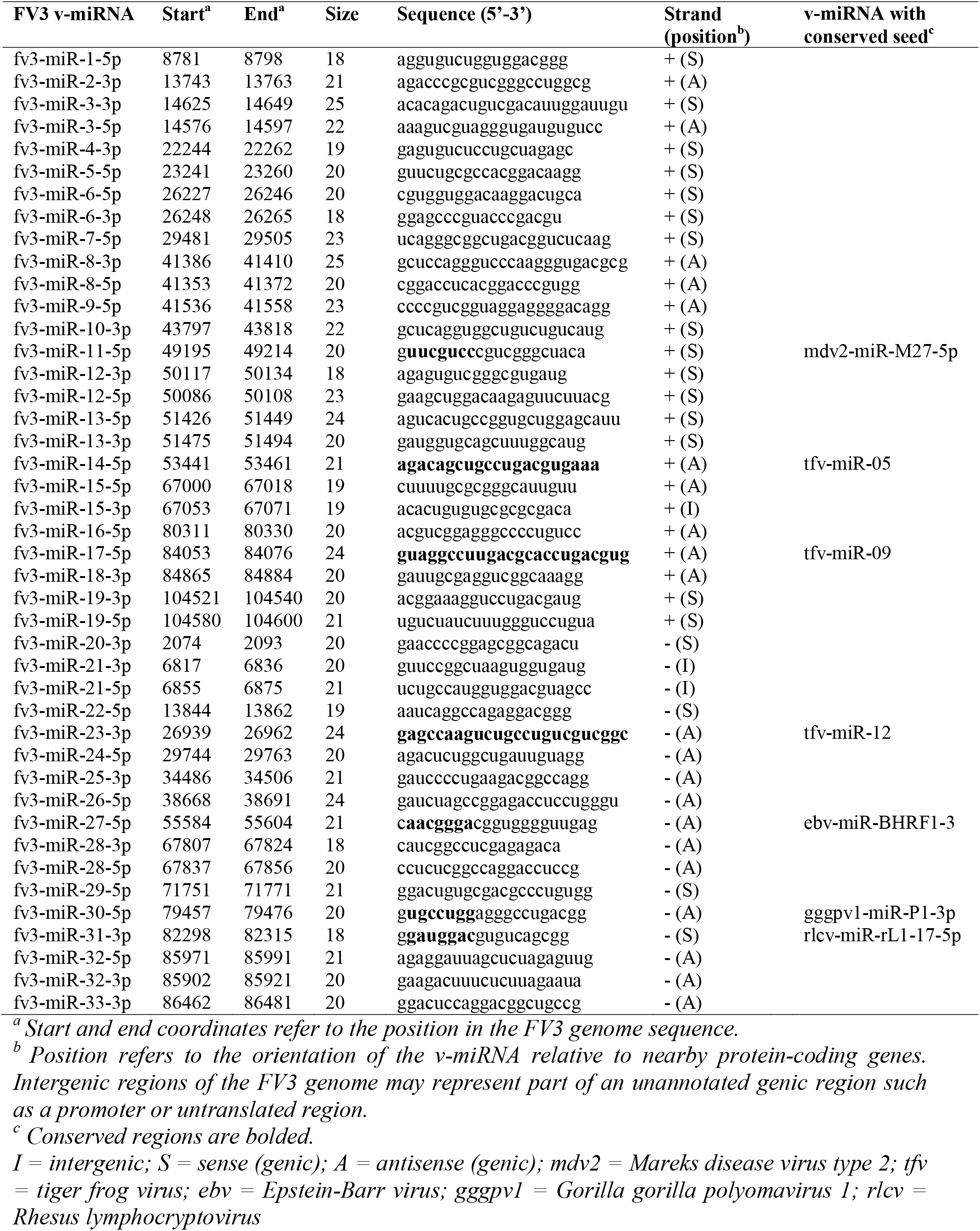
FV3 v-miRNA characteristics.

### FV3-encoded v-miRNAs are differentially expressed at different times post-infection

To gain insight into the differences in v-miRNA expression at different times post-FV3 infection, we performed differential expression analysis to compare levels of FV3 v-miRNAs at 24 hpi and 72 hpi. Read counts were normalized to their respective library size, defined as the total number of FV3-mapped miRNA reads in a given library. Differential expression analyses were performed using DESeq2 (Love et al., 2014) and EdgeR (Robinson et al., 2010). Of the 43 novel FV3 v-miRNAs, 33 were differentially expressed between the 24 hpi and 72 hpi time points (**Figure 3A**). Fifteen FV3 v-miRNAs exhibited elevated levels at 24 hpi, 18 FV3 v-miRNAs exhibited elevated levels at 72 hpi, and the levels of 10 FV3 v-miRNAs remained unchanged across the time points examined (**Figure 3A**). The FV3 v-miRNAs most strongly downregulated at 72 hpi include fv3-miR-19-5p, fv3-miR-8-5p, and fv3-miR-32-5p, while the FV3 v-miRNAs most strongly upregulated at 72 hpi include fv3-miR-30-5p, fv3-miR-8-3p, and fv3-miR-19-3p (**Figure 3A**). Clustering of the v-miRNAs according to the timing expression of the viral ORFs at the same locus (immediate early, delayed early, late, and unknown) (Majji et al., 2009) reveals that regulation of v-miRNA expression is not obviously correlated with regulation of expression of the protein-coding genes at that locus (**Figure 3B**).

**Figure 3.**
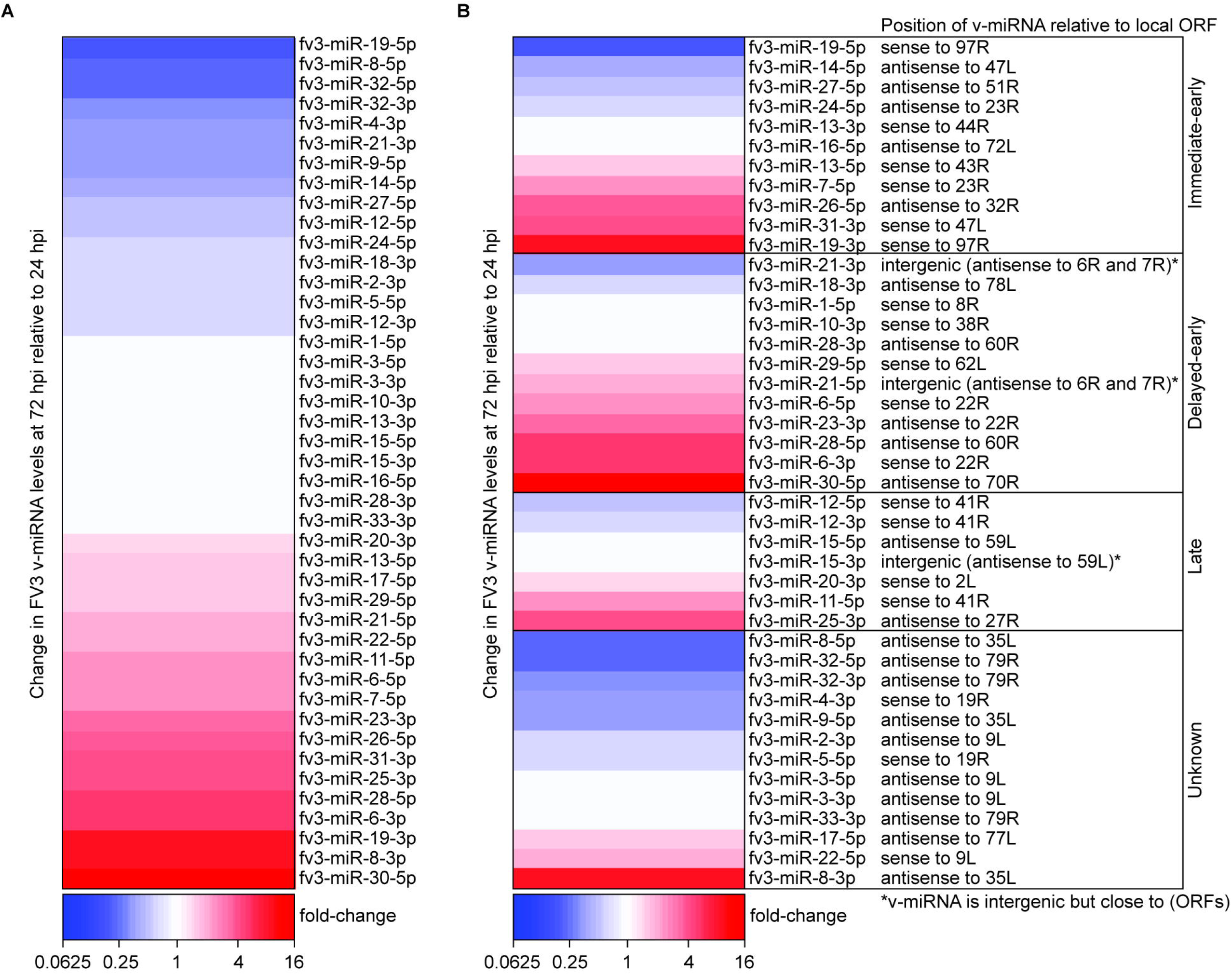
Differential expression analysis of FV3 v-miRNAs in FV3-infected Xela DS2. (**A**) Differential expression analysis was performed to compare FV3 v-miRNA levels in FV3-infected Xela DS2 cells at 72 hpi relative to 24 hpi. Statistically significant (FDR < 0.05, *n* = 3, Wald test) increases and decreases in FV3 v-miRNA levels at 72 hpi relative to 24 hpi are represented in red and blue, respectively, and white bars indicate no statistical significance. The fold-change values were generated by DESeq2 and supported by EdgeR. (**B**) The results from panel (A) clustered in relation to the timing of expression of the FV3 genes at the same locus. hpi, hours post infection; ORF, open reading frame.

### FV3-encoded v-miRNAs are predicted to target viral genes

Possible viral targets of FV3 v-miRNAs were identified by target prediction analysis using RNAhybrid (Rehmsmeier et al., 2004). Of the 43 FV3 v-miRNAs identified, 26 v-miRNAs were predicted to target 21 endogenous viral genes in 28 interactions (**Table 3** and **Table S4**). The majority of these interactions occur *in cis* (e.g. FV3 v-miRNA maps antisense to the target gene), while seven putative viral gene targets are predicted to interact with FV3 v-miRNAs *in trans* (i.e. FV3 v-miRNAs are not transcribed directly opposite the target gene) (**Table 3**). With two exceptions (fv3-miR-28-3p and fv3-miR-33-3p), FV3 v-miRNAs each appear to target a single viral gene. However, some FV3 genes (e.g. ORFs 9L, 35L, 60R, 79R, and 97R) are predicted targets of multiple FV3 v-miRNAs. FV3 v-miRNAs are predicted to target a variety of immediate early, delayed early, and late genes, as well as several genes of unknown timing of expression (**Table 3**). Timing of FV3 v-miRNA expression is not obviously related to the timing of expression of the viral target gene (**Table 3**). Most FV3 genes are still categorized as hypothetical, thus we are unable to determine how the v-miRNAs that target these genes may function to regulate the viral replicative cycle. However, examination of the characterized FV3 gene targets indicates that some FV3 v-miRNAs target viral genes that are thought to be important to the host-pathogen interaction. For example, fv3-miR-25-3p targets the FV3 tyrosine kinase (ORF27R), which contains domains that are conserved in *Xenopus* IRF proteins suggesting it may be able to mimic host IRF proteins to interfere with IFN signaling (Tian et al., 2021a).

**Table 3.**
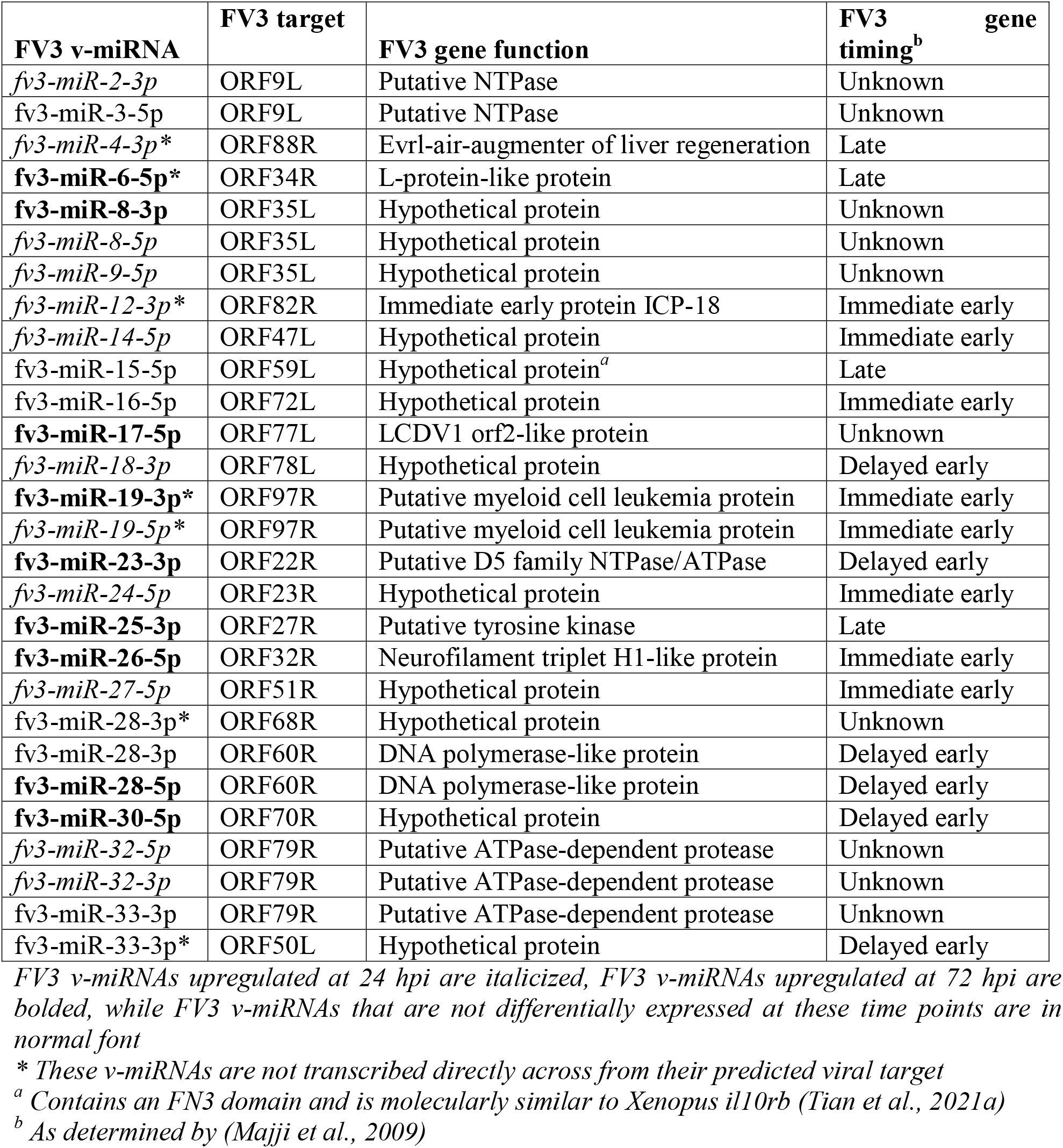
FV3 gene targets of FV3 v-miRNAs.

### FV3-encoded v-miRNAs are predicted to target host immune-related genes

Additional target prediction analyses revealed that the 43 FV3 v-miRNAs are predicted to target 1,822 *X. laevis* host genes in 1990 interactions (**Table S5**). While most of these interactions are with genes of known function, 41 FV3 v-miRNAs are predicted to target 313 *X. laevis* genes with currently unknown or unannotated functions (323 interactions). Of note, numerous FV3 v-miRNAs are predicted to target host immune-related genes such as interferon receptors (*ifnar2* and *ifnlr1*), interleukins and their receptors (*il12b, il17re, il5ra, il6r*, and *il7r*), interleukin/IFN regulatory factors (*irak1* and *irf10*), NF-κB-related factors (*nfkbia/ikka*), TGF-β-related factors (*tgfb2* and *tgfbr3*), TNF-related factors (*tnfaip8, tnfsf11, tnfsf15, traf1*, and *traf6*), genes involved in cytosolic viral nucleic acid sensing pathways (*trim25, sting1, dhx33*, and *atrx*), and even endogenous host genes involved in microRNA biogenesis/function (*drosha, ago1*, and *tarbp2*) (**Table 4**).

**Table 4.** A subset of predicted *X. laevis* immune-related gene targets of select FV3 v-miRNAs.

| **FV3 v-miRNA** | **Key *X. laevis* immune target(s)** |
| --- | --- |
| fv3-miR-1-5p | *sting1, usp4* |
| fv3-miR-4-3p | *smurf1, trim25* |
| fv3-miR-5-5p | *dhx33, tarbp2* |
| fv3-miR-6-5p | *traf6* |
| fv3-miR-6-3p | *tp53, usp38* |
| fv3-miR-7-5p | *prkcd* |
| fv3-miR-8-3p | *ifnar2, nfkbia/ikka* |
| fv3-miR-8-5p | *ifnlr1, trim7* |
| fv3-miR-9-5p | *fadd* |
| fv3-miR-10-3p | *smurf1* |
| fv3-miR-11-5p | *fadd, tgfb2* |
| fv3-miR-14-5p | *tak1* |
| fv3-miR-15-5p | *trim11* |
| fv3-miR-15-3p | *il7r, isg20l2* |
| fv3-miR-17-5p | *atrx* |
| fv3-miR-18-3p | *isg20l2* |
| fv3-miR-19-3p | *tgfbr3, usp25* |
| fv3-miR-19-5p | *tak1* |
| fv3-miR-20-3p | *il6r, irf10* |
| fv3-miR-22-5p | *ago1, atrx* |
| fv3-miR-24-5p | *irak1* |
| fv3-miR-25-3p | *nmi* |
| fv3-miR-27-5p | *atrx, prkcd, usp25* |
| fv3-miR-28-5p | *il17re, il5ra, prkcd, trim25* |
| fv3-miR-30-5p | *card9, il12b* |
| fv3-miR-31-3p | *drosha* |
| fv3-miR-33-3p | *stim1, trim7, usp5* |

To achieve an overall view of antiviral signaling pathways targeted by FV3 v-miRNAs, we mapped FV3 v-miRNAs onto pathway schematics depicting cytosolic (cGAS-STING, RIG-I/MDA5) and membrane/endosomal (TLR3, TLR4, TLR9) pattern recognition receptors (PRRs) that may be involved in sensing FV3 and the downstream antiviral pathways they activate that contribute to IRF and NF-κB transcription factor activation and the expression of pro-inflammatory cytokines and type I IFNs (**Figure 4**). FV3 is a dsDNA virus that undergoes DNA replication in two stages, first in the nucleus and then in the cytosol (Goorha, 1982; Goorha et al., 1978). The cGAS-STING pathway senses viral dsDNA in the cytosol and culminates in the production of type I IFNs (reviewed in (Hopfner and Hornung, 2020)). cGAS binding to dsDNA activates the transmembrane protein STING1, which subsequently activates the kinase TBK1. TBK1 complexes with the IKK kinase, which activates IRF3 and IRF7 transcription factors via phosphorylation to induce the transcription of type I IFNs. Additional regulators include STIM1, a calcium sensor that inhibits STING1 (Srikanth et al., 2019), the TBK1-inhibiting E3-ubiquitin ligase TRIM11 and de-ubiquitinating enzyme USP38 (Lee et al., 2013; Lin et al., 2016), FADD, an adaptor protein that is required for efficient induction of IRF7 (Balachandran et al., 2007), and the STAT-interacting protein NMI, which is a repressor of IRF7 (Wang et al., 2013a). FV3 v-miRNAs were predicted to target *sting1* (fv3-miR-1-5p), *ikka* (fv3-miR-8-3p), *stim1* (fv3-miR-33-3p), *fadd* (fv3-miR-9-5p and fv3-miR-11-5p), *trim11* (fv3-miR-15-5p), and *usp38* (fv3-miR-6-3p) (**Figure 4**). These FV3 v-miRNAs can be grouped based on predicted functional outcomes into repression of positive regulators of cGAS-STING (*sting1, ikka*, and *fadd*) or inhibition of repressors of cGAS-STING (*stim1, trim11*, and *usp38*).

**Figure 4.**
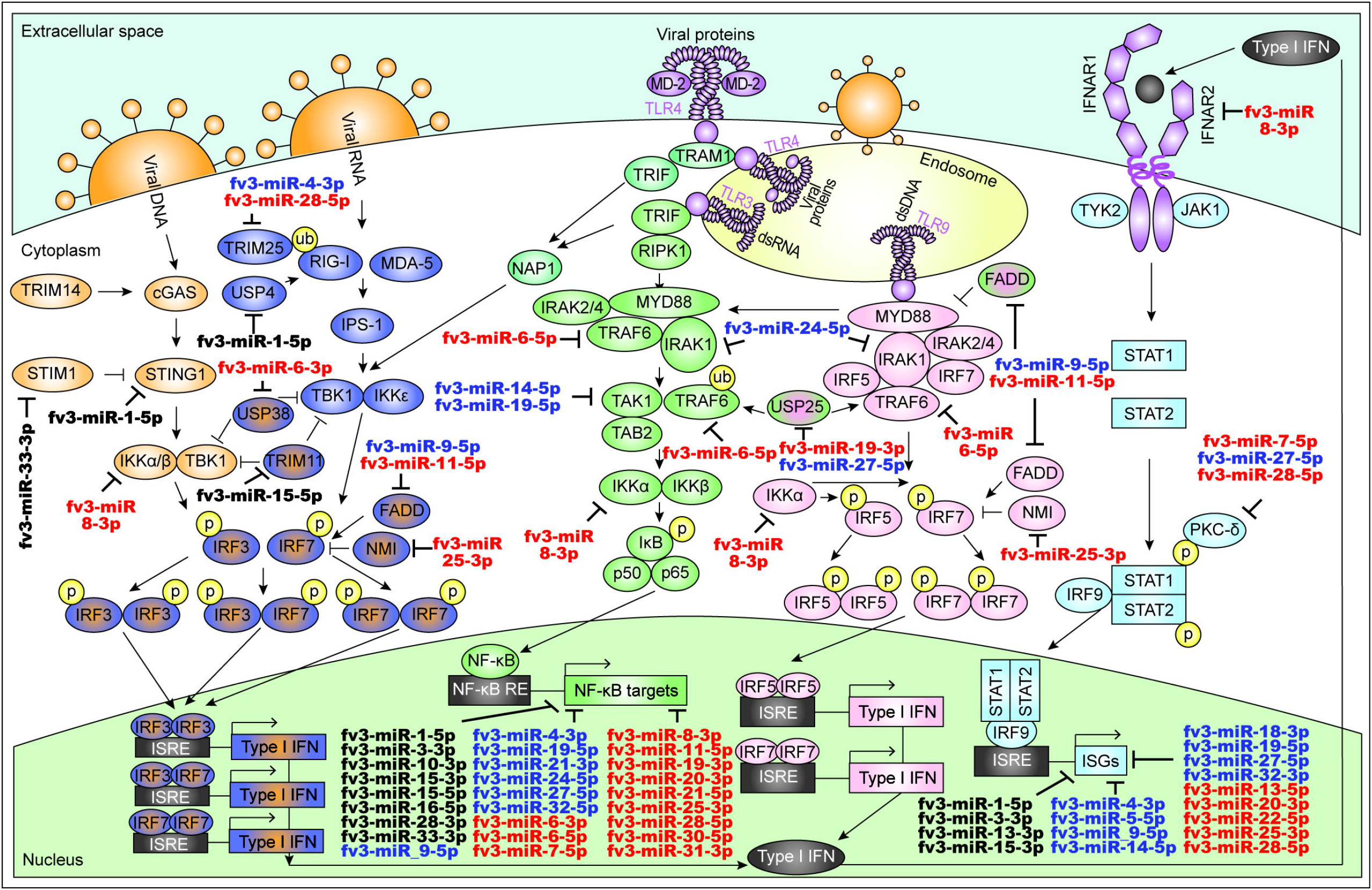
Key antiviral pathways targeted by FV3 v-miRNAs. A schematic representation of antiviral signaling pathways and the FV3 v-miRNAs that target key genes in these pathways. FV3 v-miRNAs that are downregulated at 72 hours post infection (hpi) relative to 24 hpi are depicted in blue, v-miRNAs that are upregulated at 72 hpi relative to 24 hpi are depicted in red, and FV3 v-miRNAs with consistent levels at 24 and 72 hpi are depicted in black. Base pathways were adapted from (InvivoGen, 2021) and (Todd et al., accepted). Arrow heads indicate activation/promotion while blunted heads indicate repression. p, phoshphorylation; ub, ubiquitination; NF-κB, nuclear factor kappa B; RE, response element; ISRE, interferon-stimulated response element; ISGs, interferon-stimulated genes; IFN, interferon; dsDNA, double-stranded DNA; dsRNA, double-stranded RNA.

Cytosolic viral dsRNA produced during FV3 replicative cycles (Doherty et al., 2016) may be sensed by the RIG-I/MDA5 pathway that senses dsRNA in the cytosol and initiates a signaling pathway that culminates in the production of type I IFNs (reviewed in (Rehwinkel and Gack, 2020)). dsRNA sensing by RIG-I/MDA5 activates TBK1 and IKK kinases, which results in the production of type I IFNs as described above. Important regulators of the RIG-I/MDA5 pathway include the E3-ubiquitin ligase TRIM25 which ubiquitinates and activates RIG-I (Gack et al., 2007) and USP4 that de-ubiquitinates and represses RIG-I (Wang et al., 2013b). FV3 v-miRNAs may impact the RIG-I/MDA5 pathway by targeting *fadd, trim11, nmi*, and *usp38*, as described above, and *trim25* (fv3-miR-4-3p and fv3-miR-28-5p) and *usp4* (fv3-miR-1-5p) (**Figure 4**). Thus, similar to the cGAS-STING pathway, FV3 v-miRNAs may inhibit RIG-I/MDA5 function by repressing factors that promote its function (*trim25* and *fadd*) or may promote the RIG-I/MDA5 pathway by inhibiting its repressors (*trim11, usp38*, and *usp4*).

Additional PRRs that may be involved in sensing FV3 are TLRs including TLR3 and TLR9 that respectively sense dsRNA and dsDNA in the endosome, and TLR4 which senses viral proteins at the cell membrane or in the endosome (reviewed in (Kawasaki and Kawai, 2014)). TLR3 signaling is initiated by the recognition of dsRNA in the endosome, culminating in the transcription of type I IFNs or NF-κB-induced genes through a TRIF-dependent pathway or the MYD88-dependent pathway, respectively (reviewed in (Kawasaki and Kawai, 2014)). The TRIF-dependent pathway feeds into the TBK1/IKK pathway described above, with FV3 v-miRNAs predicted to target *usp38* (fv3-miR-6-3p), *trim11* (fv3-miR-15-5p), and *fadd* (fv3-miR-9-5p and fv3-miR-11-5p). With regards to the MYD88 pathway, TLR3 ligation induces the formation of the MYD88 complex (containing MYD88 and the IRAK1, IRAK2 and IRAK4 kinases) and its interaction with the TRAF6 adaptor, which activates the TAK1 kinase. TAK1 activation activates the IKK kinase, resulting in the phosphorylation and degradation of IκB and translocation of NF-κB into the nucleus to induce transcription of NF-κB-regulated genes. FV3 v-miRNAs were predicted to target *irak1* (fv3-miR-24-5p), *traf6* (fv3-miR-6-5p), the TRAF6-stabilizing de-ubiquitinating enzyme *usp25* (fv3-miR-19-3p and fv3-miR-27-5p) (Lin et al., 2015), the repressor of MYD88 *fadd* (fv3-miR-9-5p and fv3-miR-11-5p) (Zhande et al., 2007), *tak1* (fv3-miR-14-5p and fv3-miR-19-5p), *ikka* (fv3-miR-8-3p), and numerous genes that are induced by NF-κB. (**Figure 4**). Thus, FV3 v-miRNAs may regulate TLR3 signaling by repressing positive regulators (*irak1, traf6, tak1*, and *ikka*) and negative regulators (*usp38, trim11, usp25* and *fadd*) of the pathway. Alternatively, FV3 may be sensed by the TLR9 pathway that senses dsDNA in the endosome. TLR9 ligation induces the formation of the MYD88 complex and its association with TRAF6, IRF5, and IRF7, which activates the IRF5 and IRF7 transcription factors (through phosphorylation by IKKα), triggering the production of type I IFNs (reviewed in (Kawasaki and Kawai, 2014)). FV3 v-miRNAs target the MYD88 complex, *usp25, fadd, ikka*, and *nmi* as described above (**Figure 4**). Thus, FV3 v-miRNAs may impact the TLR9 pathway by inhibiting positive regulators (*irak1, traf6, tak1, ikka*) and negative regulators (*usp25, fadd*, and *nmi*) of TLR9 signaling. Lastly, the TLR4 pathway could be initiated by sensing FV3 viral proteins in the extracellular space or the endosome (reviewed in (Kawasaki and Kawai, 2014)). FV3 v-miRNAs were not predicted to target additional genes involved in the TLR4 pathway other than *usp38, trim11*, and *fadd* as described above (**Figure 4**).

All the sensing pathways described above that may be involved in responding to FV3 infection can result in the production of type I IFNs. FV3 v-miRNAs were also predicted to target genes involved in the type I IFN signaling pathway, in which sensing of extracellular type I IFN initiates a signaling pathway that results in the transcription of thousands of ISGs (reviewed in (Platanias, 2005)). Type I IFN is recognized by the IFNAR1 and IFNAR2 receptors, which activates the TYK2 and JAK1 kinases that phosphorylate STAT1 and STAT2. STAT1 phosphorylation by PKCδ is also required for its activation (Uddin et al., 2002). STAT1/2 phosphorylation drives the formation of a STAT1-STAT2-IRF9 complex that stimulates the transcription of ISGs. FV3 v-miRNAs were predicted to target *ifnar2* (fv3-miR-8-3p), *prkcd* (fv3-miR-7-5p, fv3-miR-27-5p, and fv3-miR-28-5p), and numerous ISGs (**Figure 4**). Thus, FV3 v-miRNAs may interfere with type I IFN signaling by repressing key IFN receptors, blocking the activation of transcription factors that drive the transcription of important ISGs, or by directly repressing ISG expression.

GO term enrichment analysis was performed to begin uncovering other cellular pathways that may be broadly targeted by FV3 v-miRNAs. The targets of FV3 v-miRNAs were enriched in 28 GO term categories with broad functions across the cell such as “cellular response to cAMP”, “regulation of protein secretion”, “protein phosphorylation”, “proteolysis”, “response to abiotic stimulus”, “positive regulation of cell communication”, and “regulation of intracellular signal transduction” (**Figure 5**). While these pathways may have implications in cellular antiviral responses, we did not detect enrichment of any immune-specific categories.

**Figure 5.**
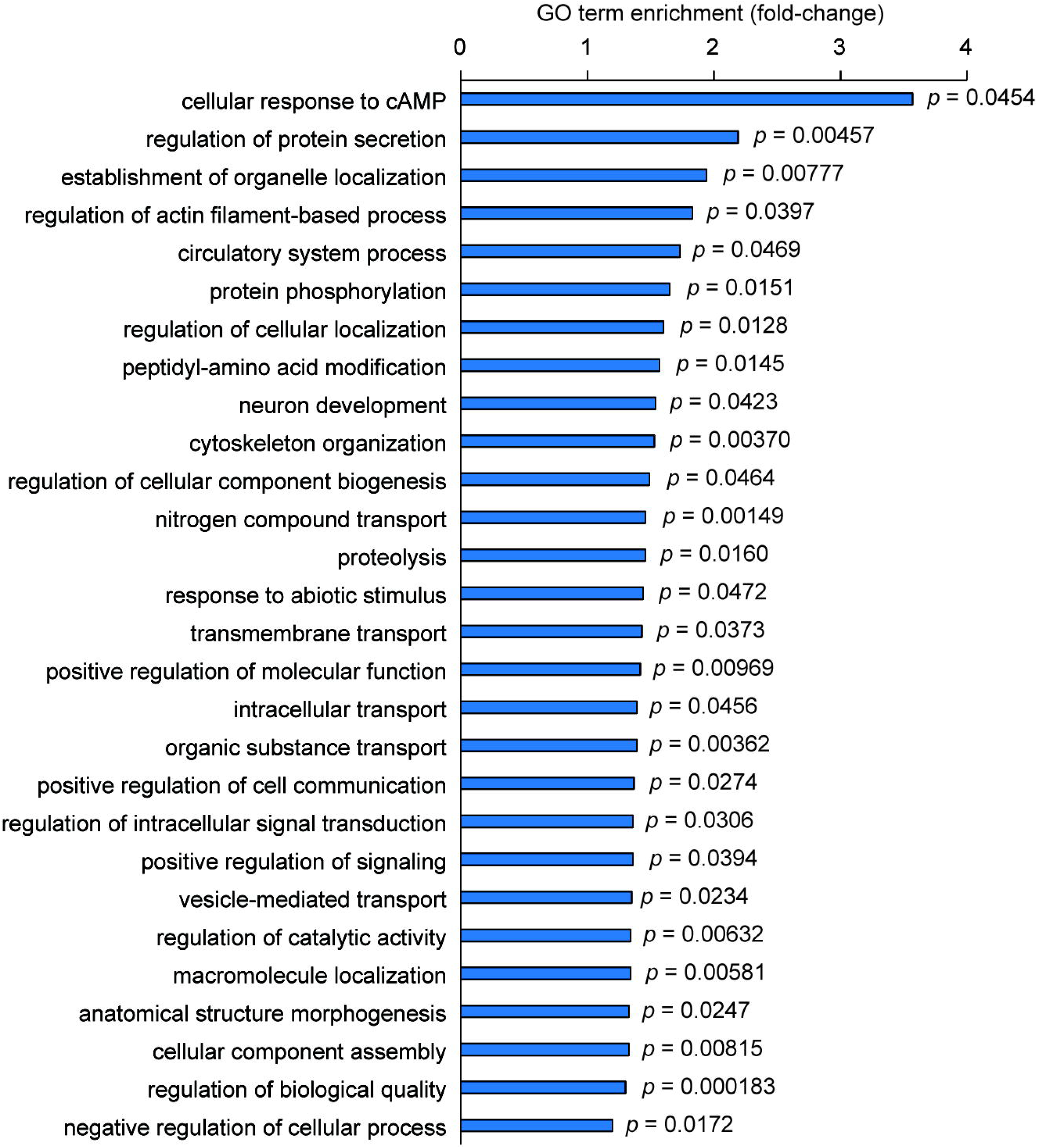
Gene ontology (GO) term analysis of host gene targets. Represented GO terms enriched in the list of *X. laevis* targets of FV3 v-miRNAs. In cases where several GO terms originating from the same hierarchy were enriched, the most specific GO term is represented here. *p*-values are indicated beside each bar and were considered statistically significant if *p* < 0.05, as determined by Fisher’s exact test. cAMP, cyclic adenosine monophosphate.

## Discussion

FV3 is immunoevasive and research in the field has centered on elucidating the involvement of FV3-encoded proteins in immunoevasion of host antiviral responses. In the past 13 years, two studies have predicted the existence of hundreds of FV3 pre-miRNAs and v-miRNAs, with each study predicting the existence of a unique set from the same genomic information. Although FV3 v-miRNAs were previously predicted to exist, the incongruence between predictions and the unusually high number of predicted v-miRNAs pose a challenge in elucidating which predictions are most accurate and which v-miRNAs, if any, are expressed in virally infected host cells. Using an experimental approach (small RNA-seq), we are the first to discover 43 mature v-miRNAs produced by FV3 during infection of a *X. laevis* dorsal skin epithelial-like cell line, representing the first report to present experimental evidence for the expression of mature FV3 v-miRNAs during cellular infection. Importantly, the small RNA library preparation chemistry used herein facilitates the discovery of transcripts that have specifically been processed into small RNAs and not degradation products of longer viral RNAs. As discussed below, the 43 FV3 v-miRNAs we discovered are unique from previous *in silico* predictions, are differentially regulated throughout infection, and target endogenous viral genes as well as a plethora of host genes, many of which are involved in antiviral immune responses. Our findings demonstrate that FV3 produces v-miRNAs that are dynamically regulated throughout infection and suggest that FV3 v-miRNAs are involved in immunoevasive strategies. *Bioinformatics pipeline validation*

We first validated the v-miRNA detection pipeline used in this study on publicly available EBV small RNA-seq data. This pipeline identified the majority of known EBV v-miRNAs but missed two and predicted a number of additional novel v-miRNAs. The lack of detection of two known EBV v-miRNAs by the pipeline is not clear, and the identification of a few novel v-miRNAs could either represent increased predictive power of the pipeline or could represent false positive predictions. However, many of the novel EBV v-miRNAs originate from the same precursor as a known v-miRNA and thus represent the other half of the miRNA:miRNA* duplex, suggesting they are likely real. Since this pipeline can accurately detect a core set of v-miRNAs, we believe this pipeline to be robust in the detection of novel FV3 v-miRNAs. Further investigation and systematic validation of each of the 43 FV3-encoded v-miRNAs discovered in this study will be an important next step in verifying the presence and functions of these v-miRNAs.

### Detection of FV3 v-miRNAs

The quantity and relative positioning of FV3 v-miRNAs discovered in this study is consistent with previous discoveries of v-miRNAs encoded by other dsDNA viruses. The discovery of 43 FV3 v-miRNAs is in line with the number of v-miRNAs identified in better-studied viruses such as EBV which encodes 44 v-miRNAs, and is within the upper threshold of the number of v-miRNAs identified in any one virus to date (70; rLCV) (Kozomara et al., 2019). The vast majority of FV3 v-miRNAs discovered herein are encoded within genic regions. v-miRNAs are encoded within genic regions in other viruses including ranaviruses, non-ranaviral iridovirids, and distantly related dsDNA viruses that infect humans. TFV, a ranavirus that infects tiger frogs (*Rana tigrina rugulosa*) and is closely related to FV3, encodes at least 24 v-miRNAs. TFV v-miRNAs are widely distributed across the genome, and all but one TFV v-miRNA are located within ORFs (Yuan et al., 2016). SGIV is a non-ranaviral iridovirid that infects fish. Sixteen SGIV v-miRNAs are scattered across the genome, representing a mix of genic (13 v-miRNAs) and intergenic (three v-miRNAs) positioning (Yan et al., 2011). v-miRNAs are also often encoded within genic regions in more distantly related dsDNA viruses, such as EBV and KSHV (Cai et al., 2005; Cai et al., 2006; Pfeffer et al., 2005). While the comparisons herein are limited to a handful of viruses as some studies have not explicitly stated v-miRNA orientations relative to protein-coding genes, it is clear that v-miRNAs are not always encoded intergenically. As viral genomes are compact, the encoding v-miRNAs within genic regions optimizes use of the genomic space and may allow for additional layers of regulation, such that transcription of a v-miRNA could occur independent of mRNA transcription (i.e. distinct promoters) or may depend on mRNA transcription (i.e. transcription of the mRNA and subsequent cleavage to produce the v-miRNA) (Marsico et al., 2013; Morales et al., 2017).

Conservation of v-miRNAs highlights their common importance in infection strategies and immunoevasion. Interestingly, three of the FV3 v-miRNAs we discovered have orthologous v-miRNAs present in TFV that share identical sequences (tfv-miR-05, tfv-miR-09, and tfv-miR-12) (Yuan et al., 2016). While the TFV host species and disease characteristics are generally distinct from FV3 and their geographical distributions only partially overlap (i.e. in China) (Allender, 2019; Weng et al., 2002), the FV3 genome is > 90% identical to that of TFV (Tan et al., 2004). Thus, it is perhaps unsurprising that the FV3 and TFV genomes encode conserved v-miRNAs. Additionally, several FV3 v-miRNAs exhibit seed sequence conservation with v-miRNAs encoded by other dsDNA viruses (MDV, EBV, gggpv1, and rLCV) (Kozomara et al., 2019) with which FV3 is more distantly related. While UTR sequences are not generally well conserved among different species, there are some cases where miRNA target sites are conserved, even in distantly related species such as frogs and humans (Todd et al., accepted). In fact, miRNA target sites are thought to be one of the most highly conserved regulatory motifs found in 3’ UTRs (Simkin et al., 2020). Thus, these v-miRNAs with shared seed sequences may represent conserved regulators of host-pathogen interactions.

v-miRNAs can be differentially expressed at different stages of infection (Broekema and Imperiale, 2013; Hussain et al., 2008; Murphy et al., 2008) and are often expressed in tissue-cell type- or host species-specific manners (Meyer et al., 2010; Yuan et al., 2016). Thus, while we detected 43 v-miRNAs in the system employed in this study, we cannot rule out the possibility that other FV3 v-miRNAs may be expressed at different time points, or in other cell types, tissues, or hosts. In this study, small RNA-seq was employed to detect and determine the exact sequence of small RNAs present in FV3-infected Xela DS2 cells, and this approach yielded 43 FV3 v-miRNAs. However, previous studies have computationally predicted v-miRNA sequences from the FV3 genome. A v-miRNA database creation study that computationally predicted pre-miRNA sequences from all viruses with genome sequences available at the time predicted 239 FV3 pre-miRNAs but did not go on to predict mature v-miRNA sequences from these pre-miRNAs (Li et al., 2008). A study published during preparation of our manuscript predicted FV3-encoded pre-miRNAs and their associated mature v-miRNAs from transcribed intergenic regions, predicting 295 pre-miRNAs that were each predicted to produce only one mature v-miRNA. Interestingly, these is no overlap in the pre-miRNAs predicted by these two studies (Li et al., 2008; Tian et al., 2021b). The much greater number of FV3 v-miRNAs detected by computational prediction analyses (Li et al., 2008; Tian et al., 2021b) – upwards of 500 – may reflect known difficulties in accurately predicting pre-miRNAs and mature v-miRNAs. Computational prediction of viral pre-miRNAs is associated with many false positives, and accurately predicting mature v-miRNA sequences from the predicted hairpins is very difficult without small RNA-seq data to guide the analysis (Demirci et al., 2017; Thomassen et al., 2006). In addition to there being no overlap between the FV3 pre-miRNAs predicted by Li et al (2008) and those predicted by Tian et al (2021b), there is also no overlap between the mature v-miRNAs predicted by Tian et al (2021b) and those detected by the small RNA-seq experiments performed herein. The latter point may be partially because the Tian et al (2021) study predicted FV3 v-miRNAs solely from intergenic regions, while our sequencing efforts suggest that FV3 v-miRNAs are largely encoded within genic regions, with only three being intergenic. Nevertheless, the possibility that the computationally predicted FV3 v-miRNAs are real cannot be ruled out, as we recognize that our study has likely not detected all FV3 v-miRNAs that exist. In addition, the analysis pipeline used herein may have underestimated the total number of v-miRNAs produced by FV3 as reads with counts < 10 per library were filtered out to improve the signal-to-noise ratio, thus FV3 v-miRNAs that are expressed at low levels may have been overlooked. Additionally, reads that mapped to the *X. laevis* genome were filtered out, as it would be impossible to identify the source (host or viral) of these miRNAs, therefore FV3 v-miRNAs with perfect complementarity to host gene transcripts may have been excluded. However, v-miRNAs do not typically exhibit perfect complementarity to host gene targets (Skalsky and Cullen, 2010). Thus, while we have discovered what is likely only a subset of the v-miRNAs produced by FV3, our study has paved the way for future studies to perform extensive v-miRNA profiling in a variety of FV3-infected cell types, tissues, and hosts.

### Impacts of FV3 v-miRNAs on viral gene expression

Target prediction analyses suggest that FV3 v-miRNAs may play a role in regulating 21 endogenous viral transcripts, including 11 characterized proteins (NTPase, Evrl-air-augmenter of liver regeneration, L-protein-like protein, ICP-18, LCDV1 orf2-like protein, myeloid cell leukemia protein, D5 family NTPase/ATPase, tyrosine kinase, neurofilament triplet H1-like protein, DNA polymerase, and ATP-dependent protease) and 10 uncharacterized (hypothetical) proteins. It is important to note that since FV3 UTRs are largely unannotated, the target prediction analyses herein focused on FV3 cds and thus might reflect an underrepresentation of the viral transcripts regulated by FV3 v-miRNAs. Nevertheless, several of the characterized FV3 gene targets regulate viral replication, indicating that FV3 v-miRNAs may impact viral replication. Indeed, v-miRNAs are known to regulate viral replication in other dsDNA viruses such as EBV, KSHV, hCMV, and Hepatitis B virus (Abend et al., 2012; Abend et al., 2010; Hancock et al., 2017; Hook et al., 2014; Landais et al., 2015; Lau et al., 2016; Lei et al., 2010). However, future functional studies are necessary to characterize the exact roles of FV3 v-miRNAs in regulating viral replication.

Of the 43 FV3 v-miRNAs discovered herein, 15 were more highly expressed at 24 hpi, 18 were more highly expressed at 72 hpi, and 10 remained at constant levels at both time points. This suggests that a subset of FV3 v-miRNAs may play important roles in the establishment of infection, while another subset might be key during the later stages of infection, and the remainder may be utilized throughout infection. This is similar to what has been previously observed, as v-miRNAs expressed during late infection have been shown to function in repressing early viral gene expression (Broekema and Imperiale, 2013; Murphy et al., 2008). For example, the insect dsDNA virus Heliothis virescens ascovirus upregulates a v-miRNA during late stages of infection to target the viral DNA polymerase and decrease viral replication (Hussain et al., 2008). Similarly, fv3-miR-28-5p is upregulated at 72 hpi, and it is predicted to target the FV3 viral DNA polymerase (ORF60R). Interestingly, the levels of FV3 v-miRNAs are not correlated with the timing of expression of the viral ORFs at the same locus, suggesting that v-miRNA expression is regulated in a manner that may be generally independent of the regulation of viral ORF expression. This is consistent with the fact that transcription of miRNAs encoded within genic regions is often driven by their own promoter (Liu et al., 2018). Additionally, the levels of FV3 v-miRNAs are not perfectly correlated with the timing of expression of the viral ORFs that they are predicted to target. These discrepancies could be because the time points examined in our study might not optimally align with the time points typically associated with early and late viral gene expression. For example, the study that determined the timing of FV3 gene expression utilized an MOI of 25 and examined viral gene expression at 2, 7, and 9 hpi, while our study infected cells at an MOI of 2 and examined v-miRNA expression at 24 and 72 hpi. Thus, future studies should examine the expression of FV3 v-miRNAs over a time course of infection with a larger number of sampling points, and perform mRNA-seq simultaneously to improve the ability to correlate FV3 v-miRNA and mRNA expression.

### Impacts of FV3 v-miRNAs on host gene expression

v-miRNAs are emerging post-transcriptional regulators of host immune responses and have been ascribed immunoevasive functions [reviewed in (Cullen, 2013)]. For example, v-miRNAs produced by other dsDNA viruses (EBV, hCMV, KSHV) have been found to target host antiviral genes resulting in repression of proinflammatory cytokine production and release (Abend et al., 2012; Abend et al., 2010; Bouvet et al., 2021; Hancock et al., 2017; Hook et al., 2014; Landais et al., 2015; Lau et al., 2016; Lei et al., 2010). Accordingly, target prediction and pathway analyses revealed that FV3 v-miRNAs are predicted to target host genes involved in several antiviral pathways such as the cGAS-STING, RIG-I/MDA5, TLR3, TLR4, TLR9, and type I IFN signaling pathways, as well as the downstream products of signaling pathways triggered by virus sensing by these PRRs (NF-κB-induced genes and ISGs), and the impacts of these potential interactions are discussed below. It is, however, important to note that while annotation of the *X. laevis* genome is improving, the tetraploid nature of *X. laevis* has proved challenging for proper scaffolding and gene annotation. As a result, *X. laevis* gene annotations are incomplete, and immune genes are particularly poorly annotated. Thus, our target prediction analyses likely reflect an underrepresentation of the host transcripts targeted by FV3 v-miRNAs. For example, the *X. laevis* genome encodes an expanded type I IFN family (Sang et al., 2016), but the UTRs of these genes are not annotated and are thus missing from our analysis.

FV3 v-miRNAs were predicted to target multiple components of the cGAS-STING, RIG-I/MDA5, TLR3, TLR4, TLR9, and type I IFN pathways. The antiviral gene targets represent a combination of regulators of the PRRs that initially sense viral macromolecules, regulators of the downstream antiviral pathways that such sensing triggers, and the products of these signaling pathways that function to establish antiviral programs. This not only suggests that FV3 may have evolved mechanisms to evade these pathways, but that these pathways are potentially important to anti-FV3 defenses, a research area that remains largely unexamined. v-miRNAs produced by FV3 are predicted to target genes that promote the function of these pathways, as well as genes that inhibit these pathways. While miRNAs function to repress their targets through degradation or translational repression, miRNAs can upregulate their targets by promoting their transcription by interacting with enhancer elements (Xiao et al., 2017), stabilizing their mRNA by blocking repressor binding sites (Eiring et al., 2010), or promoting their translation through various mechanisms (Mengardi et al., 2017; Vasudevan et al., 2007). This may explain why FV3 v-miRNAs are predicted to target activators and repressors of the same pathway. Thus, the mechanisms through which miRNAs impact gene expression are complex, and it is therefore essential to systematically analyze the function of each FV3 v-miRNA discovered herein.

Several candidate FV3 v-miRNAs will be particularly exciting to functionally characterize. For example, the IRAK1 kinase involved in TLR signaling is targeted by fv3-miR-24-5p, and the human IRAK1 ortholog is a known target of a KHSV v-miRNA (Abend et al., 2012). In addition, it is interesting to note that two FV3 v-miRNAs upregulated at different times during infection (fv3-miR-9-5p and fv3-miR-11-5p) target FADD, which is important to the regulation of cGAS-STING, RIG-I/MDA5, TLR3, TLR4, and TLR9 signaling pathways. This suggests that FV3 has evolved the ability to simultaneously impact at least five different antiviral signaling pathways that defend against viruses at the cell membrane, in the cytoplasm, and in the endosome. As impairment of type I IFN signaling has been observed in other viruses such as EBV, hCMV, and KSHV (Abend et al., 2012; Bouvet et al., 2021; Hooykaas et al., 2017; Huang and Lin, 2014; Huang et al., 2015), it will be important to functionally characterize fv3-miR-8-3p, fv3-miR-7-5p, fv3-miR-27-5p, and fv3-miR-28-5p which target mediators of type I IFN signaling. It is also interesting to note that FV3 v-miRNAs appear to mostly target the RIG-I/MDA5 cytoplasmic sensing pathway by specifically targeting genes involved in ubiquitination (*trim25, usp4, usp38*, and *trim11*), suggesting that FV3 may largely regulate this pathway by modulating ubiquitination. Lastly, while FV3 v-miRNAs that are differentially expressed across the time points examined appear to target the antiviral pathways examined in roughly equal proportions, FV3 v-miRNAs that are sustained in their levels across the time points examined (fv3-miR-33-3p, fv3-miR-1-5p, and fv3-miR-15-5p) specifically target the cGAS-STING and RIG-I/MDA5 cytosolic viral nucleic acid sensing pathways and do not appear to target any membrane/endosomal pathways (TLR3, TLR4, and TLR9) or type I IFN signaling. This suggests that FV3 may produce these v-miRNAs at sustained levels throughout infection to continually evade the cGAS-STING and RIG-I/MDA5 sensing systems, which is logical as FV3 produces both dsDNA and dsRNA during its replication in the cytoplasm, which is where these sensing systems operate. While speculation into the immunoevasive functions of FV3 v-miRNAs is an important first step, follow-up studies are needed to validate the predicted interactions between FV3 v-miRNAs and host antiviral genes. It will also be important to manipulate FV3 v-miRNA expression and assess the impacts on these host antiviral pathways and viral replication to reveal the potential immunoevasive functions of FV3 v-miRNAs.

v-miRNAs sometimes function as mimics of host miRNAs (Kincaid et al., 2012), and bioinformatics analyses suggest that up to 26% of known v-miRNAs produced by viruses that afflict humans could mimic their human host miRNAs as they contain identical seed sequences (Kincaid and Sullivan, 2012). In a previous study, we examined changes in the expression of host miRNAs in response to FV3 infection and, interestingly, FV3 v-miRNAs target the same antiviral pathways as *X. laevis* host miRNAs that are differentially expressed in response to FV3 infection (Todd et al., accepted). However, despite the overlap in pathways targeted, FV3 v-miRNAs and host miRNAs appear to target generally distinct sets of genes in these pathways within the *in vitro* infection model used in this study. Thus, the present data suggests that it is unlikely that the FV3 v-miRNAs we detected function as mimics of host miRNAs in this system, as FV3 v-miRNA targets are generally distinct from host miRNA targets, and FV3 v-miRNAs do not exhibit any seed sequence conservation with *X. laevis* miRNAs. Characterization of FV3 v-miRNAs expressed under different conditions is needed before the possibility that FV3 v-miRNAs can function as host miRNA mimics can be ruled out.

GO term analysis was used to uncover the cellular pathways targeted by FV3 v-miRNAs. Analysis of the predicted host gene targets of FV3 v-miRNAs revealed the statistical enrichment of terms related to antiviral signaling such as “regulation of intracellular signal transduction”, “regulation of protein secretion”, “protein phosphorylation”, and “positive regulation of cell communication”. The most enriched category is “cellular response to cAMP”, which is interesting as cAMP levels have been linked to IFN activity (Weber and Stewart, 1975), cAMP-dependent activation of protein kinase A is known to modulate the transcription of many cytokines (e.g. *tnf, il6*, and *il10*) (Johansson et al., 2004) and has been shown to attenuate virus-induced disruption of epithelial barriers (Rezaee et al., 2017). Thus, if cellular responses to cAMP play a role in the frog skin epithelial response to FV3, FV3 may modulate such responses as a mechanism of immunoevasion. Interestingly, immune-specific pathways are not overrepresented in the list of host gene targets of FV3 v-miRNAs, but rather FV3 v-miRNAs appear to target a wide range of cellular processes. However, it is important to note that the lack of complete annotations for many *X. laevis* immune genes may have impacted the ability to predict immune-related gene targets, which may have contributed to the lack of immune pathway enrichment identified by GO analysis. Nevertheless, FV3 v-miRNAs may impact antiviral responses by broadly impacting host cell function rather than focusing their regulation on immune-specific signaling pathways alone.

Cells can broadly regulate their function through modulating miRNA biogenesis and function. Interestingly, viruses are known alter global host gene expression by interfering with host miRNA pathways (Andersson et al., 2005; Lu and Cullen, 2004). Viral targeting of host miRNA pathways can be beneficial to viruses as it allows them to manipulate the expression of numerous genes simultaneously, including important antiviral genes that must be activated to promote effective immune responses. Several FV3 v-miRNAs are predicted to target key components of miRNA pathways including *drosha1, ago1*, and *tarbp2.* By targeting these genes, FV3 may be able to inhibit pre-miRNA processing (via *drosha1*) as well as RISC assembly and function (via *ago1* and *tarbp2).* Therefore, dampening of host miRNA biogenesis and function may be a conserved mechanism of viral immunoevasion employed by FV3.

## Conclusions

Our study challenges the theory that ranaviral proteins are the major players in mediating immunoevasion by discovering that viral non-coding RNAs such as v-miRNAs may play key roles in disrupting host antiviral programs. The findings herein advance our understanding of ranavirus host-pathogen interactions and add to a growing body of literature that links v-miRNAs to immunoevasion. While characterization of the FV3 v-miRNAs detected herein is anticipated to provide immense functional insight, this likely only touches the surface of the numerous mechanisms through which viruses such as FV3 manipulate their hosts. Our attempts, along with the attempts of others to uncover the functions of viral non-coding RNAs are helping to pivot the ranavirus field into a new era of research.

## Acknowledgements

We thank Maxwell P. Bui-Marinos for their technical assistance in setting up the FV3 infections.

## Supplementary material

**Figure S1. Confirmation of viral gene transcription in FV3-infected Xela DS2 cells.** RT-PCR was used to detect immediate early (viral caspase activation and recruitment domain-containing protein, *vCARD*), delayed early (viral polymerase II*α vPOLIIα*), and late (major capsid protein, *MCP*) FV3 gene transcripts in cDNA generated from uninfected and FV3-infected Xela DS2 cells at 24- and 72-hours post infection (hpi). Detection of the cel-miR-39 spike-in served as a positive control for miRNA detection, and *X. laevis* beta-actin (*actb*) was detected as a positive control to ensure amplifiability. NTC, no-template control. Results from three independent experiments (*n* = 3) are shown.

**Table S1.** Primers used in this study.

**Table S2.** Validation of v-miRNA detection pipeline on publicly available EBV small RNA-seq data.

**Table S3.** FV3 pre-miRNA characteristics.

**Table S4.** Predicted interactions between FV3 v-miRNAs and FV3 cds.

**Table S5.** Predicted interactions between FV3 v-miRNAs and *X. laevis* 3’ UTRs.

## References

Abend, J.R., Ramalingam, D., Kieffer-Kwon, P., Uldrick, T.S., Yarchoan, R., Ziegelbauer, J.M., 2012. Kaposi’s sarcoma-associated herpesvirus microRNAs target IRAK1 and MYD88, two components of the toll-Like Receptor/interleukin-1R signaling cascade, to reduce inflammatory-cytokine expression. J Virol 86, 11663–11674.

Abend, J.R., Uldrick, T., Ziegelbauer, J.M., 2010. Regulation of tumor necrosis factor-Like weak Inducer of apoptosis receptor protein (TWEAKR) expression by Kaposi’s sarcoma-associated herpesvirus microRNA Prevents TWEAK-induced apoptosis and inflammatory cytokine expression. J Virol 84, 12139–12151.

Afgan, E., Baker, D., van den Beek, M., Blankenberg, D., Bouvier, D., Čech, M., Chilton, J., Clements, D., Coraor, N., Eberhard, C., Grüning, B., Guerler, A., Hillman-Jackson, J., Von Kuster, G., Rasche, E., Soranzo, N., Turaga, N., Taylor, J., Nekrutenko, A., Goecks, J., 2016. The Galaxy platform for accessible, reproducible and collaborative biomedical analyses: 2016 update. Nucleic Acids Res 44, W3–W10.

Allender, M.C., 2019. Ranaviral disease in reptiles and amphibians, Fowler’s Zoo and Wild Animal Medicine Current Therapy, Volume 9, pp. 364–370.

Andersson, M.G., Haasnoot, P.C.J., Xu, N., Berenjian, S., Berkhout, B., Akusja□rvi, G.r., 2005. Suppression of RNA interference by adenovirus virus-associated RNA. J Virol 79, 9556–9565.

Andrews, S., 2010. FastQC: A quality control tool for high throughput sequence data [online].

Aranda, P.S., LaJoie, D.M., Jorcyk, C.L., 2012. Bleach gel: A simple agarose gel for analyzing RNA quality. Electrophoresis 33, 366–369.

Ashburner, M., Ball, C.A., Blake, J.A., Botstein, D., Butler, H., Cherry, J.M., Davis, A.P., Dolinski, K., Dwight, S.S., Eppig, J.T., Harris, M.A., Hill, D.P., Issel-Tarver, L., Kasarskis, A., Lewis, S., Matese, J.C., Richardson, J.E., Ringwald, M., Rubin, G.M., Sherlock, G., 2000. Gene ontology: Tool for the unification of biology. The Gene Ontology Consortium. Nat Genet 25, 25–29.

Balachandran, S., Venkataraman, T., Fisher, P.B., Barber, G.N., 2007. Fas-associated death domain-containing protein-mediated antiviral innate immune signaling involves the regulation of Irf7. J Immunol 178, 2429–2439.

Besch, R., Poeck, H., Hohenauer, T., Senft, D., Häcker, G., Berking, C., Hornung, V., Endres, S., Ruzicka, T., Rothenfusser, S., Hartmann, G., 2009. Proapoptotic signaling induced by RIG-I and MDA-5 results in type I interferon-independent apoptosis in human melanoma cells. J Clin Invest 119, 2399–2411.

Bouvet, M., Voigt, S., Tagawa, T., Albanese, M., Chen, Y.-F.A., Chen, Y., Fachko, D.N., Pich, D., Göbel, C., Skalsky, R.L., Hammerschmidt, W., Miller, M.S., 2021. Multiple viral microRNAs regulate interferon release and signaling early during infection with Epstein-Barr Virus. mBio 12.

Brand, M.D., Hill, R.D., Brenes, R., Chaney, J.C., Wilkes, R.P., Grayfer, L., Miller, D.L., Gray, M.J., 2016. Water temperature affects susceptibility to ranavirus. Ecohealth 13, 350–359.

Breuer, K., Foroushani, A.K., Laird, M.R., Chen, C., Sribnaia, A., Lo, R., Winsor, G.L., Hancock, R.E., Brinkman, F.S., Lynn, D.J., 2013. InnateDB: Systems biology of innate immunity and beyond--recent updates and continuing curation. Nucleic Acids Res 41, D1228–1233.

Broekema, N.M., Imperiale, M.J., 2013. miRNA regulation of BK polyomavirus replication during early infection. Proc Natl Acad Sci U S A 110, 8200–8205.

Bui-Marinos, M.P., Todd, L.A., Wasson, M.D., Morningstar, B.E.E., Katzenback, B.A., 2021. Prior induction of cellular antiviral pathways limits frog virus 3 replication in two permissive *Xenopus laevis* skin epithelial-like cell lines. Dev Comp Immunol 124, 104200.

Bui-Marinos, M.P., Varga, J.F.A., Vo, N.T.K., Bols, N.C., Katzenback, B.A., 2020. Xela DS2 and Xela VS2: Two novel skin epithelial-like cell lines from adult African clawed frog (*Xenopus laevis*) and their response to an extracellular viral dsRNA analogue. Dev Comp Immunol 112.

Cai, X., Lu, S., Zhang, Z., Gonzalez, C.M., Damania, B., Cullen, B.R., 2005. Kaposi’s sarcoma-associated herpesvirus expresses an array of viral microRNAs in latently infected cells. Proc Natl Acad Sci U S A 102, 5570–5575.

Cai, X., Schafer, A., Lu, S., Bilello, J.P., Desrosiers, R.C., Edwards, R., Raab-Traub, N., Cullen, B.R., 2006. Epstein-Barr virus microRNAs are evolutionarily conserved and differentially expressed. PLoS Pathog 2, e23.

Carey, C., Cohen, N., Rollins-Smith, L., 1999. Amphibian declines: An immunological perspective. Dev Comp Immunol 23, 459–472.

Chen, G., Ward, B.M., Yu, K.H., Chinchar, V.G., Robert, J., 2011. Improved knockout methodology reveals that frog virus 3 mutants lacking either the 18K immediate-early gene or the truncated vIF-2alpha gene are defective for replication and growth *in vivo*. J Virol 85, 11131–11138.

Chinchar, V.G., Hyatt, A., Miyazaki, T., Williams, T., 2009. Family Iridoviridae: poor viral relations no longer, Lesser Known Large dsDNA Viruses, pp. 123–170.

Cullen, B.R., 2013. MicroRNAs as mediators of viral evasion of the immune system. Nat Immunol 14, 205–210.

Daszak, P., Berger, L., Cunningham, A.A., Hyatt, A.D., Green, D.E., Speare, R., 1999. Emerging infectious diseases and amphibian population declines. Emerg Infect Dis 5, 735–748.

De Jesus Andino, F., Grayfer, L., Chen, G., Chinchar, V.G., Edholm, E.S., Robert, J., 2015. Characterization of Frog Virus 3 knockout mutants lacking putative virulence genes. Virology 485, 162–170.

Demirci, M.D.S., Baumbach, J., Allmer, J., 2017. On the performance of pre-microRNA detection algorithms. Nat Commun 8.

Doherty, L., Poynter, S.J., Aloufi, A., DeWitte-Orr, S.J., 2016. Fish viruses make dsRNA in fish cells: characterization of dsRNA production in rainbow trout *(Oncorhynchus mykiss)* cells infected with viral haemorrhagic septicaemia virus, chum salmon reovirus and frog virus 3. J Fish Dis 39, 1133–1137.

Duan, F., Liao, J., Huang, Q., Nie, Y., Wu, K., 2012. HSV-1 miR-H6 inhibits HSV-1 replication and IL-6 expression in human corneal epithelial cells *in vitro*. Clin Dev Immunol 2012, 192791.

Eiring, A.M., Harb, J.G., Neviani, P., Garton, C., Oaks, J.J., Spizzo, R., Liu, S., Schwind, S., Santhanam, R., Hickey, C.J., Becker, H., Chandler, J.C., Andino, R., Cortes, J., Hokland, P., Huettner, C.S., Bhatia, R., Roy, D.C., Liebhaber, S.A., Caligiuri, M.A., Marcucci, G., Garzon, R., Croce, C.M., Calin, G.A., Perrotti, D., 2010. miR-328 functions as an RNA decoy to modulate hnRNP E2 regulation of mRNA translation in leukemic blasts. Cell 140, 652–665.

Enk, J., Levi, A., Weisblum, Y., Yamin, R., Charpak-Amikam, Y., Wolf, D.G., Mandelboim, O., 2016. HSV1 MicroRNA Modulation of GPI Anchoring and Downstream Immune Evasion. Cell Rep 17, 949–956.

Friedländer, M.R., Mackowiak, S.D., Li, N., Chen, W., Rajewsky, N., 2012. miRDeep2 accurately identifies known and hundreds of novel microRNA genes in seven animal clades. Nucleic Acids Res 40, 37–52.

Gack, M.U., Shin, Y.C., Joo, C.-H., Urano, T., Liang, C., Sun, L., Takeuchi, O., Akira, S., Chen, Z., Inoue, S., Jung, J.U., 2007. TRIM25 RING-finger E3 ubiquitin ligase is essential for RIG-I-mediated antiviral activity. Nature 446, 916–920.

Gantress, J., Maniero, G.D., Cohen, N., Robert, J., 2003. Development and characterization of a model system to study amphibian immune responses to iridoviruses. Virology 311, 254–262.

Goorha, R., 1982. Frog virus 3 DNA replication occurs in two stages. J Virol 43, 519–528.

Goorha, R., Granoff, A., 1974. Macromolecular synthesis in cells infected by frog virus 3. I. Virus-specific protein synthesis and its regulation. Virology 60, 237–250.

Goorha, R., Murti, G., Granoff, A., Tirey, R., 1978. Macromolecular synthesis in cells infected by frog virus 3. VIII. The nucleus is a site of frog virus 3 DNA and RNA synthesis. Virology 84, 32–50.

Grayfer, L., Edholm, E., De Jesus Andino, F., Chinchar, V., Robert, J., 2015. Ranavirus host immunity and immune evasion, in: Gray, M., Chinchar, V. (Eds.), Ranaviruses. Springer, Cham.

Grey, F., Meyers, H., White, E.A., Spector, D.H., Nelson, J., 2007. A human cytomegalovirus-encoded microRNA regulates expression of multiple viral genes involved in replication. PLoS Pathog 3, e163.

Grey, F., Tirabassi, R., Meyers, H., Wu, G., McWeeney, S., Hook, L., Nelson, J.A., 2010. A viral microRNA down-regulates multiple cell cycle genes through mRNA 5′UTRs. PLoS Pathog 6, e1000967.

Hancock, M.H., Hook, L.M., Mitchell, J., Nelson, J.A., Moscona, A., Renne, R., Yurochko, A., 2017. Human cytomegalovirus microRNAs miR-US5-1 and miR-UL112-3p block proinflammatory cytokine production in response to NF-κB-activating factors through direct downregulation of IKKα and IKKβ. mBio 8.

Hook, Lauren M., Grey, F., Grabski, R., Tirabassi, R., Doyle, T., Hancock, M., Landais, I., Jeng, S., McWeeney, S., Britt, W., Nelson, Jay A., 2014. Cytomegalovirus miRNAs Target Secretory Pathway Genes to Facilitate Formation of the Virion Assembly Compartment and Reduce Cytokine Secretion. Cell Host Microbe 15, 363–373.

Hooykaas, M.J., Kruse, E., Wiertz, E.J., Lebbink, R.J., 2016. Comprehensive profiling of functional Epstein-Barr virus miRNA expression in human cell lines. BMC Genomics 17, 644.

Hooykaas, M.J.G., van Gent, M., Soppe, J.A., Kruse, E., Boer, I.G.J., van Leenen, D., Groot Koerkamp, M.J.A., Holstege, F.C.P., Ressing, M.E., Wiertz, E.J.H.J., Lebbink, R.J., 2017. EBV microRNA BART16 suppresses type I IFN signaling. J Immunol 198, 4062–4073.

Hopfner, K.-P., Hornung, V., 2020. Molecular mechanisms and cellular functions of cGAS-STING signalling. Nat Rev Mol Cell Biol 21, 501–521.

Hoverman, J.T., Gray, M.J., Miller, D.L., 2010. Anuran susceptibilities to ranaviruses: role of species identity, exposure route, and a novel virus isolate. Dis Aquat Organ 89, 97–107.

Huang, W.-T., Lin, C.-W., 2014. EBV-encoded miR-BART20-5p and miR-BART8 inhibit the IFN-γ-STAT1 pathway associated with disease progression in nasal NK-cell lymphoma. Am J Pathol 184, 1185–1197.

Huang, Y., Chen, D., He, J., Cai, J., Shen, K., Liu, X., Yang, X., Xu, L., 2015. Hcmv-miR-UL112 attenuates NK cell activity by inhibition type I interferon secretion. Immunol Lett 163, 151–156.

Huang, Y., Huang, X., Liu, H., Gong, J., Ouyang, Z., Cui, H., Cao, J., Zhao, Y., Wang, X., Jiang, Y., Qin, Q., 2009. Complete sequence determination of a novel reptile iridovirus isolated from soft-shelled turtle and evolutionary analysis of *Iridoviridae*. BMC Genomics 10, 224.

Hussain, M., Taft, R.J., Asgari, S., 2008. An insect virus-encoded microRNA regulates viral replication. J Virol 82, 9164–9170.

InvivoGen, 2021. www.invivogen.com/review-type1-ifn-production.

Jacques, R., Edholm, E.S., Jazz, S., Odalys, T.L., Francisco, J.A., 2017. Xenopus-FV3 host-pathogen interactions and immune evasion. Virology 511, 309–319.

Johansson, C.C., Bryn, T., Yndestad, A., Eiken, H.G., Bjerkeli, V., Froland, S.S., Aukrust, P., Tasken, K., 2004. Cytokine networks are pre-activated in T cells from HIV-infected patients on HAART and are under the control of cAMP. AIDS 18, 171–179.

Kärber, G., 1931. Beitrag zur kollektiven Behandlung pharmakologischer Reihenversuche. Naunyn-Schmiedebergs Archiv für Experimentelle Pathologie und Pharmakologie 162, 480–483.

Kawasaki, T., Kawai, T., 2014. Toll-like receptor signaling pathways. Front Immunol 5.

Kincaid, R.P., Burke, J.M., Sullivan, C.S., 2012. RNA virus microRNA that mimics a B-cell oncomiR. Proc Natl Acad Sci U S A 109, 3077–3082.

Kincaid, R.P., Sullivan, C.S., 2012. Virus-encoded microRNAs: An overview and a look to the future. PLoS Pathog 8.

Knudson, D.L., Tinsley, T.W., 1974. Replication of a nuclear polyhedrosis virus in a continuous cell culture of *Spodoptera frugiperda:* purification, assay of infectivity, and growth characteristics of the virus. J Virol 14, 934–944.

Kozomara, A., Birgaoanu, M., Griffiths-Jones, S., 2019. miRBase: From microRNA sequences to function. Nucleic Acids Res 47, D155–D162.

Krieg, A.M., 2002. CpG motifs in bacterial DNA and their immune effects. Annu Rev Immunol 20, 709–760.

Krug, L.T., Pozharskaya, V.P., Yu, Y., Inoue, N., Offermann, M.K., 2004. Inhibition of infection and replication of human herpesvirus 8 in microvascular endothelial cells by alpha interferon and phosphonoformic acid. J Virol 78, 8359–8371.

Landais, I., Pelton, C., Streblow, D., DeFilippis, V., McWeeney, S., Nelson, J.A., 2015. Human ctomegalovirus miR-UL112-3p trgets TLR2 and mdulates the TLR2/IRAK1/NFκB sgnaling pthway. PLOS Pathog 11.

Langland, J.O., Jacobs, B.L., 2002. The role of the PKR-inhibitory genes, E3L and K3L, in determining vaccinia virus host range. Virology 299, 133–141.

Langland, J.O., Kash, J.C., Carter, V., Thomas, M.J., Katze, M.G., Jacobs, B.L., 2006. Suppression of proinflammatory signal transduction and gene expression by the dual nucleic acid binding domains of the vaccinia virus E3L proteins. J Virol 80, 10083–10095.

Langmead, B., Salzberg, S.L., 2012. Fast gapped-read alignment with Bowtie 2. Nat Methods 9, 357–359.

Lau, B., Poole, E., Krishna, B., Montanuy, I., Wills, M.R., Murphy, E., Sinclair, J., 2016. The expression of human cytomegalovirus microRNA MiR-UL148D during latent infection in primary myeloid cells inhibits activin A-triggered secretion of IL-6. Sci Rep 6.

Law, C.W., Alhamdoosh, M., Su, S., Dong, X., Tian, L., Smyth, G.K., Ritchie, M.E., 2016. RNA-seq analysis is easy as 1-2-3 with limma, Glimma and edgeR. F1000Res 5.

Lee, Y., Song, B., Park, C., Kwon, K.-S., 2013. TRIM11 negatively regulates IFNß production and antiviral activity by targeting TBK1. PLoS One 8.

Lei, X., Bai, Z., Ye, F., Xie, J., Kim, C.-G., Huang, Y., Gao, S.-J., 2010. Regulation of NF-κB inhibitor IκBα and viral replication by a KSHV microRNA. Nat Cell Biol 12, 193–199.

Lei, X., Zhu, Y., Jones, T., Bai, Z., Huang, Y., Gao, S.J., 2012. A Kaposi’s sarcoma-associated herpesvirus microRNA and its variants target the transforming growth factor ß pathway to promote cell survival. J Virol 86, 11698–11711.

Li, S.C., Shiau, C.K., Lin, W.C., 2008. Vir-Mir db: Prediction of viral microRNA candidate hairpins. Nucleic Acids Res 36, D184–189.

Lin, D., Zhang, M., Zhang, M.X., Ren, Y., Jin, J., Zhao, Q., Pan, Z., Wu, M., Shu, H.B., Dong, C., Zhong, B., 2015. Induction of USP25 by viral infection promotes innate antiviral responses by mediating the stabilization of TRAF3 and TRAF6. Proc Natl Acad Sci U S A 112, 11324–11329.

Lin, H.R., Ganem, D., 2011. Viral microRNA target allows insight into the role of translation in governing microRNA target accessibility. Proc Natl Acad Sci U S A 108, 5148–5153.

Lin, M., Zhao, Z., Yang, Z., Meng, Q., Tan, P., Xie, W., Qin, Y., Wang, R.-F., Cui, J., 2016. USP38 inhibits type I interferon signaling by editing TBK1 ubiquitination through NLRP4 signalosome. Mol Cell 64, 267–281.

Lin, X., Liang, D., He, Z., Deng, Q., Robertson, E.S., Lan, K., 2011. miR-K12-7-5p encoded by Kaposi’s sarcoma-associated herpesvirus stabilizes the latent state by targeting viral ORF50/RTA. PLoS One 6, e16224.

Liu, B., Shyr, Y., Cai, J., Liu, Q., 2018. Interplay between miRNAs and host genes and their role in cancer. Brief Funct Genomics 18, 255–266.

Lo, A.K., To, K.F., Lo, K.W., Lung, R.W., Hui, J.W., Liao, G., Hayward, S.D., 2007. Modulation of LMP1 protein expression by EBV-encoded microRNAs. Proc Natl Acad Sci U S A 104, 16164–16169.

Lorenz, R., Bernhart, S.H., Höner Zu Siederdissen, C., Tafer, H., Flamm, C., Stadler, P.F., Hofacker, I.L., 2011. ViennaRNA Package 2.0. Algorithms Mol Biol 6, 26.

Love, M.I., Huber, W., Anders, S., 2014. Moderated estimation of fold change and dispersion for RNA-seq data with DESeq2. Genome Biol 15, 550.

Lu, S., Cullen, B.R., 2004. Adenovirus VA1 noncoding RNA can inhibit small interfering RNA and microRNA biogenesis. J Virol 78, 12868–12876.

Lu, Y., Qin, Z., Wang, J., Zheng, X., Lu, J., Zhang, X., Wei, L., Peng, Q., Zheng, Y., Ou, C., Ye, Q., Xiong, W., Li, G., Fu, Y., Yan, Q., Ma, J., 2017. Epstein-Barr virus miR-BART6-3p inhibits the RIG-I pathway. J Innate Immun 9, 574–586.

Maes, R., Granoff, A., 1967. Viruses and renal carcinoma of *Rana pipiens*. Virology 33, 491–502.

Majji, S., Thodima, V., Sample, R., Whitley, D., Deng, Y., Mao, J., Chinchar, V.G., 2009. Transcriptome analysis of Frog virus 3, the type species of the genus *Ranavirus*, family *Iridoviridae*. Virology 391, 293–303.

Marsico, A., Huska, M.R., Lasserre, J., Hu, H., Vucicevic, D., Musahl, A., Orom, U., Vingron, M., 2013. PROmiRNA: A new miRNA promoter recognition method uncovers the complex regulation of intronic miRNAs. Genome Biol 14, R84.

Martin, M., 2011. Cutadapt removes adapter sequences from high-throughput sequencing reads. EMBnet.journal 17, 10–12.

McKenzie, C.M., Piczak, M.L., Snyman, H.N., Joseph, T., Theijin, C., Chow-Fraser, P., Jardine, C.M., 2019. First report of ranavirus mortality in a common snapping turtle *Chelydra serpentina*. Dis Aquat Organ 132, 221–227.

Mengardi, C., Limousin, T., Ricci, E.P., Soto-Rifo, R., Decimo, D., Ohlmann, T., 2017. microRNAs stimulate translation initiation mediated by HCV-like IRESes. Nucleic Acids Res 45, 4810–4824.

Meyer, C., Grey, F., Kreklywich, C.N., Andoh, T.F., Tirabassi, R.S., Orloff, S.L., Streblow, D.N., 2010. Cytomegalovirus microRNA expression is tissue specific and is associated with persistence. J Virol 85, 378–389.

Meylan, E., Curran, J., Hofmann, K., Moradpour, D., Binder, M., Bartenschlager, R., Tschopp, J., 2005. Cardif is an adaptor protein in the RIG-I antiviral pathway and is targeted by hepatitis C virus. Nature 437, 1167–1172.

Mi, H., Muruganujan, A., Ebert, D., Huang, X., Thomas, P.D., 2019. PANTHER version 14: more genomes, a new PANTHER GO-slim and improvements in enrichment analysis tools. Nucleic Acids Res 47, D419–D426.

Miller, D., Gray, M., Storfer, A., 2011. Ecopathology of ranaviruses infecting amphibians. Viruses 3, 2351–2373.

Morales, S., Monzo, M., Navarro, A., 2017. Epigenetic regulation mechanisms of microRNA expression. Biomol Concepts 8, 203–212.

Murphy, E., Vanicek, J., Robins, H., Shenk, T., Levine, A.J., 2008. Suppression of immediate-early viral gene expression by herpesvirus-coded microRNAs: Implications for latency. Proc Natl Acad Sci U S A 105, 5453–5458.

Peace, A., O’Regan, S.M., Spatz, J.A., Reilly, P.N., Hill, R.D., Carter, E.D., Wilkes, R.P., Waltzek, T.B., Miller, D.L., Gray, M.J., 2019. A highly invasive chimeric ranavirus can decimate tadpole populations rapidly through multiple transmission pathways. Ecol Modell 410.

Pfeffer, S., Sewer, A., Lagos-Quintana, M., Sheridan, R., Sander, C., Grasser, F.A., van Dyk, L.F., Ho, C.K., Shuman, S., Chien, M., Russo, J.J., Ju, J., Randall, G., Lindenbach, B.D., Rice, C.M., Simon, V., Ho, D.D., Zavolan, M., Tuschl, T., 2005. Identification of microRNAs of the herpesvirus family. Nat Methods 2, 269–276.

Pham, P.H., Jung, J., Bols, N.C., 2011. Using 96-well tissue culture polystyrene plates and a fluorescence plate reader as tools to study the survival and inactivation of viruses on surfaces. Cytotechnology 63, 385–397.

Platanias, L.C., 2005. Mechanisms of type-I- and type-II-interferon-mediated signalling. Nat Rev Immunol 5, 375–386.

Price, Stephen J., Garner, Trenton W.J., Nichols, Richard A., Balloux, F., Ayres, C., Mora-Cabello de Alba, A., Bosch, J., 2014. Collapse of amphibian communities due to an introduced ranavirus. Curr Biol 24, 2586–2591.

Rehmsmeier, M., Steffen, P., Hochsmann, M., Giegerich, R., 2004. Fast and effective prediction of microRNA/target duplexes. RNA 10, 1507–1517.

Rehwinkel, J., Gack, M.U., 2020. RIG-I-like receptors: their regulation and roles in RNA sensing. Nat Rev Immunol 20, 537–551.

Rezaee, F., Harford, T.J., Linfield, D.T., Altawallbeh, G., Midura, R.J., Ivanov, A.I., Piedimonte, G., 2017. cAMP-dependent activation of protein kinase A attenuates respiratory syncytial virus-induced human airway epithelial barrier disruption. PLoS One 12, e0181876.

Robinson, M.D., McCarthy, D.J., Smyth, G.K., 2010. edgeR: A Bioconductor package for differential expression analysis of digital gene expression data. Bioinformatics 26, 139–140.

Sang, Y., Liu, Q., Lee, J., Ma, W., McVey, D.S., Blecha, F., 2016. Expansion of amphibian intronless interferons revises the paradigm for interferon evolution and functional diversity. Sci Rep 6.

Scheel, T.K.H., Moore, M.J., Luna, J.M., Nishiuchi, E., Fak, J., Darnell, R.B., Rice, C.M., 2017. Global mapping of miRNA-target interactions in cattle *(Bos taurus)*. Sci Rep 7, 8190.

Simkin, A., Geissler, R., McIntyre, A.B.R., Grimson, A., 2020. Evolutionary dynamics of microRNA target sites across vertebrate evolution. PLOS Genet 16.

Skalsky, R.L., Cullen, B.R., 2010. Viruses, microRNAs, and host interactions. Annu Rev Microbiol 64, 123–141.

Sorel, O., Dewals, B.G., 2016. MicroRNAs in large herpesvirus DNA genomes: recent advances. Biomol Concepts 7, 229–239.

Srikanth, S., Woo, J.S., Wu, B., El-Sherbiny, Y.M., Leung, J., Chupradit, K., Rice, L., Seo, G.J., Calmettes, G., Ramakrishna, C., Cantin, E., An, D.S., Sun, R., Wu, T.T., Jung, J.U., Savic, S., Gwack, Y., 2019. The Ca(2+) sensor STIM1 regulates the type I interferon response by retaining the signaling adaptor STING at the endoplasmic reticulum. Nat Immunol 20, 152–162.

Stern-Ginossar, N., Elefant, N., Zimmermann, A., Wolf, D.G., Saleh, N., Biton, M., Horwitz, E., Prokocimer, Z., Prichard, M., Hahn, G., Goldman-Wohl, D., Greenfield, C., Yagel, S., Hengel, H., Altuvia, Y., Margalit, H., Mandelboim, O., 2007. Host immune system gene targeting by a viral miRNA. Science 317, 376–381.

Sun, W., Huang, Y., Zhao, Z., Gui, J., Zhang, Q., 2006. Characterization of the Rana grylio virus 3beta-hydroxysteroid dehydrogenase and its novel role in suppressing virus-induced cytopathic effect. Biochem Biophys Res Commun 351, 44–50.

Tan, W.G., Barkman, T.J., Gregory Chinchar, V., Essani, K., 2004. Comparative genomic analyses of frog virus 3, type species of the genus *Ranavirus* (family *Iridoviridae*). Virology 323, 70–84.

The Gene Ontology Consortium, 2019. The Gene Ontology Resource: 20 years and still GOing strong. Nucleic Acids Res 47, D330–D338.

Thomassen, G.O., Rosok, O., Rognes, T., 2006. Computational prediction of microRNAs encoded in viral and other genomes. J Biomed Biotechnol 2006, 95270.

Tian, Y., De Jesús Andino, F., Khwatenge, C.N., Li, J., Robert, J., Sang, Y., 2021a. Virus-targeted transcriptomic snalyses implicate ranaviral interaction with host interferon response in frog virus 3-infected frog tissues. Viruses 13.

Tian, Y., Khwatenge, C.N., Li, J., De Jesus Andino, F., Robert, J., Sang, Y., 2021b. Targeted transcriptomics of frog virus 3 in infected frog tissues reveal non-coding regulatory elements and microRNAs in the ranaviral genome and their potential interaction with host immune response. Front Immunol 12.

Uddin, S., Sassano, A., Deb, D.K., Verma, A., Majchrzak, B., Rahman, A., Malik, A.B., Fish, E.N., Platanias, L.C., 2002. Protein kinase C-delta (PKC-delta) is activated by type I interferons and mediates phosphorylation of Stat1 on serine 727. J Biol Chem 277, 14408–14416.

Vasudevan, S., Tong, Y., Steitz, J.A., 2007. Switching from repression to activation: microRNAs can up-regulate translation. Science 318, 1931–1934.

Wang, J., Yang, B., Hu, Y., Zheng, Y., Zhou, H., Wang, Y., Ma, Y., Mao, K., Yang, L., Lin, G., Ji, Y., Wu, X., Sun, B., 2013a. Negative regulation of Nmi on virus-triggered type I IFN production by targeting IRF7. J Immunol 191, 3393–3399.

Wang, L., Zhao, W., Zhang, M., Wang, P., Zhao, K., Zhao, X., Yang, S., Gao, C., 2013b. USP4 positively regulates RIG-I-mediated antiviral response through deubiquitination and stabilization of RIG-I. J Virol 87, 4507–4515.

Wang, M., Yu, F., Wu, W., Wang, Y., Ding, H., Qian, L., 2018. Epstein-Barr virus-encoded microRNAs as regulators in host immune responses. Int J Biol Sci 14, 565–576.

Weber, J.M., Stewart, R.B., 1975. Cyclic AMP potentiation of interferon antiviral activity and effect of interferon on cellular cyclic AMP levels. J Gen Virol 28, 363–372.

Weng, S.P., He, J.G., Wang, X.H., Lu, L., Deng, M., Chan, S.M., 2002. Outbreaks of an iridovirus disease in cultured tiger frog, *Rana tigrina rugulosa*, in southern China. J Fish Dis 25, 423–427.

Winton, J., Batts, W., deKinkelin, P., LeBerre, M., Bremont, M., Fijan, N., 2010. Current lineages of the *epithelioma papulosum cyprini* (EPC) cell line are contaminated with fathead minnow, Pimephales promelas, cells. J Fish Dis 33, 701–704.

Xiao, M., Li, J., Li, W., Wang, Y., Wu, F., Xi, Y., Zhang, L., Ding, C., Luo, H., Li, Y., Peng, L., Zhao, L., Peng, S., Xiao, Y., Dong, S., Cao, J., Yu, W., 2017. MicroRNAs activate gene transcription epigenetically as an enhancer trigger. RNA Biol 14, 1326–1334.

Yan, M., He, J., Zhu, W., Zhang, J., Xia, Q., Weng, S., Xu, X., 2016. A microRNA from infectious spleen and kidney necrosis virus modulates expression of the virus-mock basement membrane component VP08R. Virology 492, 32–37.

Yan, Y., Cui, H., Jiang, S., Huang, Y., Huang, X., Wei, S., Xu, W., Qin, Q., 2011. Identification of a novel marine fish virus, Singapore grouper iridovirus-encoded microRNAs expressed in grouper cells by Solexa sequencing. PLoS One 6, e19148.

Yan, Y., Guo, C., Ni, S., Wei, J., Li, P., Wei, S., Cui, H., Qin, Q., 2015. Singapore grouper iridovirus (SGIV) encoded SGIV-miR-13 attenuates viral infection via modulating major capsid protein expression. Virus Res 205, 45–53.

Yuan, J.M., Chen, Y.S., He, J., Weng, S.P., Guo, C.J., He, J.G., 2016. Identification and differential expression analysis of MicroRNAs encoded by Tiger Frog Virus in cross-species infection *in vitro*. Virol J 13, 73.

Zhande, R., Dauphinee, S.M., Thomas, J.A., Yamamoto, M., Akira, S., Karsan, A., 2007. FADD negatively regulates lipopolysaccharide signaling by impairing interleukin-1 receptor-associated kinase 1-MyD88 interaction. Mol Cell Biol 27, 7394–7404.

Zhang, B.C., Zhang, J., Sun, L., 2014. In-depth profiling and analysis of host and viral microRNAs in Japanese flounder *(Paralichthys olivaceus)* infected with megalocytivirus reveal involvement of microRNAs in host-virus interaction in teleost fish. BMC Genomics 15, 878.

