## Supplementary figures and images for "Discovery of frog virus 3 microRNAs and their roles in evasion of host antiviral responses"

### Supplementary Figure 1

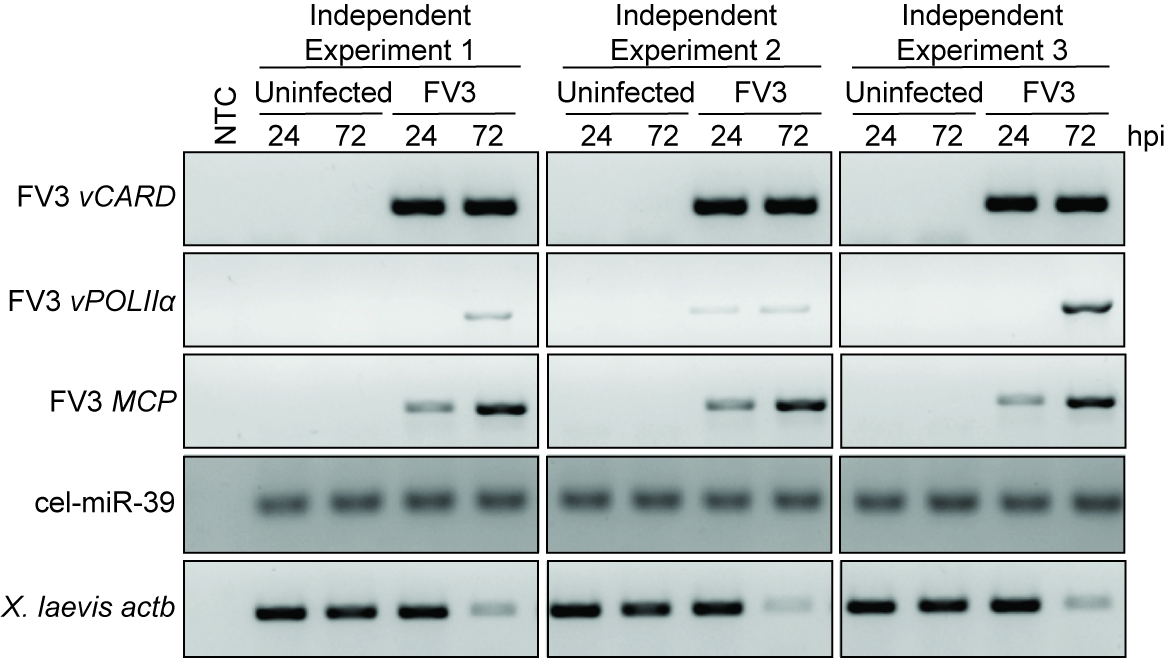
